# A single cell atlas defines perinatal factors that drive mouse bone marrow development

**DOI:** 10.64898/2026.01.21.700817

**Authors:** Brian M Dulmovits, Carson Shalaby, Fangfang Song, James Garifallou, Joshua Bertels, Fanxin Long, Christopher S Thom

## Abstract

Processes that direct initial colonization and maturation of the bone marrow remain elusive, despite lifelong importance to hematopoiesis. Bone marrow mesenchymal and stromal cell (BMSC) maturation establishes supportive niches prior to hematopoietic stem cell (HSC) recruitment. We define mouse BMSC progenitor identities and temporal emergence in a developmental single cell atlas spanning fetal life through 18 months of age. We clarify Cxcl12-abundant reticular (CAR) cell and osteoblast development, including temporal emergence of signaling modalities that direct HSC quiescence and regenerative capacity. CAR cells are absent until birth, with a developmental block during fetal maturation related to transcriptional changes during perinatal life. Temporal changes in *Early B Cell Factor (Ebf) 1-3* expression and activity correlate with CAR formation and niche establishment, including Ebf2 repression and induction of Ebf3 activities that direct metabolic changes. Changes in systemic physiology underlie these transcriptional changes, including perinatal induction of lipid metabolism, inflammation, and hypoxic signaling. These systemic factors direct CAR cell emergence after birth, providing temporal and developmental resolution underlying the colonization and maturation of the bone marrow environment.

## Introduction

Bone marrow hematopoiesis supports lifelong blood cell production. Hematopoietic niches in bone marrow use direct cell-cell communication, extracellular matrix components, and secreted signals to support hematopoietic stem and progenitor cell (HSPC) cell retention, quiescence, and lineage potential. These niches are comprised of bone marrow stromal cells (BMSCs), including Leptin receptor (Lepr)^+^ CXC chemokine ligand 12 (Cxcl12)-Abundant Reticular (CAR) cells, as well as Nestin-expressing mesenchymal and endothelial cells, microvascular endothelial cells, specialized megakaryocytes, endosteal osteoblasts, and other cell types.^1–7^ These cells produce CXCL12, stem cell factor (SCF, Kitl), pleiotrophin (PTN), and insulin-like growth factor 1 (IGF1).^8–11^ Early B-cell Factor 3 (*Ebf3*) or Forkhead Box C1 (*Foxc1*) transcription factor activities are crucial for LepR^+^ CAR cell formation and function.^12,13^ However, the cellular and transcriptional regulatory events that spur niche formation and function are not fully understood.

The bone marrow does not support hematopoiesis until the late second trimester in humans and postnatally in mice.^14^ An absence of HSCs for much of mouse fetal life results from a lack of hematopoietic niche stroma, which arise in late gestation and mature through early postnatal life.^9,15,16^ The colonization of long-term HSCs in the bone marrow coincides with Lepr^+^ BMSC emergence after birth, suggesting that niche development in the neonatal period is essential to bone marrow maturation.

The developmental origins, patterning, and transcriptional control of BMSC emergence are critical to understanding physiologic and molecular determinants that construct the hematopoietic niche. *Cxcl12*, *Nestin*, and *Lepr* reporter mice have been used to map adult niche cell responses during steady-state, stress, or marrow injury, and fate mapping with *Prrx1* and *Col2a1* constructs have elucidated cellular origins of some adult marrow stromal populations.^17–19^ Yet single cell transcriptomics approaches have revealed marked transcriptional heterogeneity in BMSC, identifying distinct adipo-osteolineage trajectories among cells with CAR-like transcriptional profiles.^20–24^ Developmental cues that regulate the formation of Lepr^+^ CAR cells are only beginning to be studied.^25^

We reasoned that a longitudinal single cell transcriptomic atlas was necessary to ascertain developmental origins for CAR cells, allowing us to define physiologic and signaling processes that drive CAR cell formation in the perinatal period. Using published and novel data sets, we provide a mechanistic framework for perinatal BMSC and CAR emergence. Functional CAR cells only emerge postnatally, consistent with signaling output and gene expression signatures that indicate HSC support only in the postnatal period. Finally, we identify developmental regulation of key CAR transcription factor activities as well as inflammatory, metabolic, and hypoxic regulatory pathways that may dictate CAR cell formation and function. These findings reveal bone marrow patterning necessary for long term hematopoiesis, based on events that occur in the perinatal period. These events fit with the dramatic physiologic changes that accompany the fetal-to-neonatal transition. Perturbations in these processes would be expected to perturb CAR cell formation and function, with adverse downstream impacts on HSC fitness.

## Results

### An integrated single cell atlas of mouse bone marrow ontogeny

We generated a single cell atlas of mouse bone marrow development comprising 19 single cell RNA-sequencing (scRNAseq) samples spanning late gestation (embryonic day 16.5, E16.5) to 18 months (18 mo) aged adults (**Fig. 1A**). This atlas incorporated published data^9,15,24,26,27^ and new scRNAseq data from postnatal day 10 (P10) and 18 mo marrow (‘Dulmovits’ data). Bone marrow cells were generally isolated from hindlimbs using a combination of purification and enzymatic digestion methods (**Supplemental Tables 1-2**).

**Figure 1.**
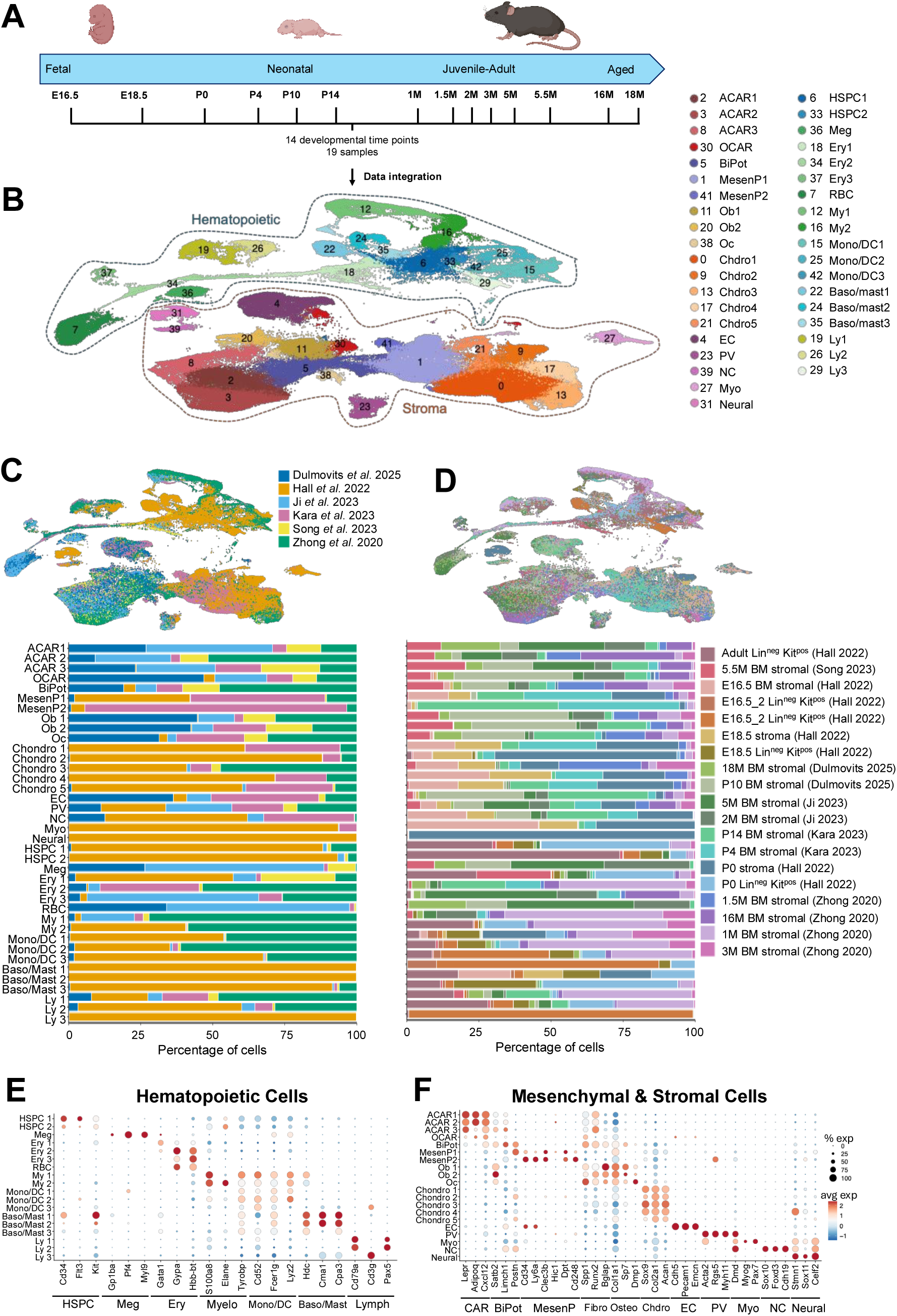
Generation of integrated single cell atlas of mouse bone marrow development. (A) Schematic of analysis and workflow using scVI integration. (B) UMAP projection of scRNAseq data from 161,339 hematopoietic and stromal cells spanning E16.5-18mo, resulting in 41 distinct clusters. Individual cells are colored by cluster. (C) UMAP projection of complete dataset with individual cells colored by original dataset and bar plot displaying the contribution of each original dataset per cluster. (D) UMAP projection of complete dataset with individual cells colored by original scRNAseq sample and bar plot displaying the contribution of each sample per cluster. (E) Dot plot of select hematopoietic marker genes. (F) Dot plot of select stromal marker genes.

Following quality control and integration (**Methods** and **Supplemental Fig. 1**), we observed excellent sample integration across data sets, with all data sources consistently contributing to most cell clusters (**Fig. 1B-D**). We ultimately found that clusters comprised predominantly from a single dataset or time point reflected true biological and/or developmental differences, rather than integration artifact.

Within these integrated data, we identified all major hematopoietic and stromal lineages within the bone marrow, based on marker gene expression (**Fig. 1B**). Cell types included hematopoietic stem and progenitor cells (*Cd34*, *Flt3*, *Kit*), erythroid (*Gypa*, *Gata1*, *Hbb-bt*), myeloid (*S100a8*, *Elane*, *Tyrobp*), monocytic-dendritic (*Cd52*, *Fcer1g*, *Lyz2)*, basophil-mast (*Hdc*, *Cma1*, *Cpa3*), lymphoid (*Cd79a*, *Cd3g*, *Pax5*) and megakaryocytic cells (*Gp1ba*, *Pf4*, *Myl9*; **Fig. 1E**). We also resolved 20 stromal cell clusters, including endothelial (*Cdh5*, *Emcn*, *Pecam*), perivascular (*Acta2*, *Rgs5*, *Myh11*), neuronal and neural crest (*Celf2*, *Stmn1*, *Sox10*), and mesenchymal progenitor cells, along with terminal lineage-differentiated cells (*Ly6a*, *Postn*, *Lepr, Acan*, *Dmp1*; **Fig. 1F**). This developmental atlas formed the basis for defining cell types and pathways that dictate bone marrow hematopoietic niche emergence throughout the mouse lifespan.

### Understanding the emergence of major mesenchymal stromal cells and the hematopoietic niche

Given our interest in CAR cell development, we focused analyses on 96,252 bone marrow mesenchymal lineage cells from 15 clusters (**Methods**, **Fig. 2A**, **Supplemental Fig. 1**). We identified five stromal cell types, including HSC-supporting CAR cells, mesenchymal stem cells (MSC), fibroblasts, osteoid lineage (*Col1a1*, *Ifitm5*, *Dmp1*) and chondrocyte lineage progenitors (*Sox9*, *Acan*, *Col2a1*; **Fig. 2B**).

**Figure 2.**
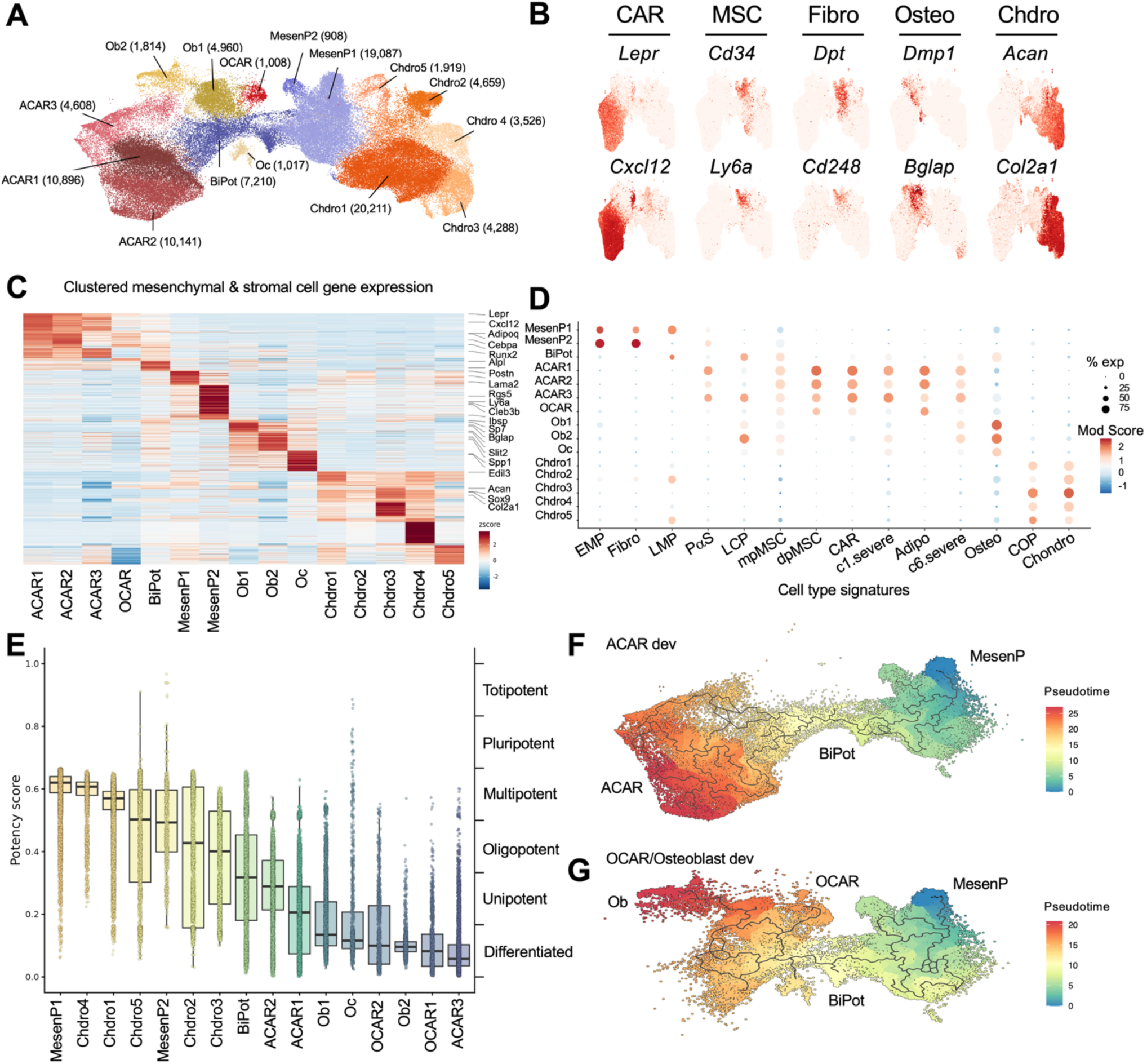
Integrated single cell atlas defines the ACAR lineage trajectory. (A) UMAP projection of scRNAseq data from 96,252 mesenchymal lineage cells spanning E16.5-18mo resulting in 15 clusters. Individual cells colored by cluster, and cell number per cluster is displayed in parentheses. (B) Feature plots of select lineage marker genes. (C) Heatmap of the 100 most differentially expressed genes per cluster. (D) Dot plot of module scores related to previously published mesenchymal progenitor and terminal populations. (E) Whisker plot of potency score calculated using CytoTRACE2. (F-G) Lineage trajectories constructed by arranging transcriptional cell states along pseudotime with Monocle3. MesenP1 cells were set as the root of the trajectory. UMAP projection with trajectory line and distance along pseudotime are displayed for (F) ACAR development (G) OCAR/osteogenic lineage development.

Putative niche stroma clusters express *Cxcl12*, *Kitl*, and *Lepr*. We therefore subdivided the CAR cell population into three adipo-CXCL12 abundant reticular cell (ACAR1-3) and one osteo-CXCL12 abundant reticular cell (OCAR) clusters based on lineage gene expression and UMAP location (**Fig. 2C**).^23^ ACAR1 and ACAR2 possessed higher expression of adipocytic markers (*Lpl*, *Adipoq*, *Cebpa*). ACAR3 contained cells with somewhat higher expression of osteoblast genes (*Runx2*, *Alpl*, *Bglap*), and pathway analysis confirmed enrichment of skeletal and limb development in the ACAR3 transcriptome (**Supplemental Fig. 2A-B**). Few cells contained markers of white, brown, or beige adipocytes, indicating that our ACAR and OCAR populations are distinct from mature adipocytes (**Supplemental Fig. 2C**). Instead, ACAR transcriptional profiles suggest dual functionality as i) hematopoietic niche stroma and ii) a stromal progenitor reservoir capable of responding to adipogenic and osteogenic stimuli.^4^

We also identified two mesenchymal progenitors subclusters (MesenP1-2) and a bipotential progenitor cluster (BiPot), named for its proximity to both the adipo- and osteo-lineage cells (**Fig. 2A-C**). MesenP cells showed high expression of mesenchymal stem cell markers (*Cd34*, *Ly6a*, *Thy1*, *Pdgfra*) and fibroblast markers (*Dpt*, *Cd248*, *Cd44*; **Fig. 2B-C**). Among MesenP cells, we observed rare *Hic1^+^* and *Pthlh^+^* cells (**Supplemental Fig. 3A**). These have been shown to represent multipotent appendicular mesenchymal progenitors and skeletal stem cells that have an ability to produce bone marrow stromal cells.^28,29^ Consistent with these expectations, MesenP1-2 cell transcriptomes matched those of early and late mesenchymal progenitor cells (EMPs, LMPs)^6^ and PDGFRα^+^ Sca-1^+^ (PaS) cells^9^ based on coordinate gene expression ‘modules’ compared with established cell types^30^ (**Fig. 2D** and **Supplemental Fig. 4**). Some MesenP cells also expressed universal fibroblast markers (**Supplemental Fig. 3B**).

We further identified two groups of BiPot cells. BiPot cells proximal to MesenP clusters expressed high levels of *Postn*, *Edil3*, *Tnn*, *Aspn*, and *Dkk3,* similar to late mesenchymal progenitor cells^24^ (LMP, **Fig. 2D**). BiPot cells that more closely resembled Osteoblast and ACAR clusters on our dimension reduction plot were marked by *Satb2*, *Slit2*, and *Wif1* (**Fig. 2D** and **Supplemental Fig. 3C-D**). This subpopulation of BiPot cells co-expressed adipocyte and osteoblast genes, suggesting that these cells represent more committed adipo-osteo-progenitors (**Fig. 2D** and **Supplemental Fig. 3C-D**).

Osteolineage and chrondrolineage cells possessed high module scores for validated osteoblast and chondrocyte gene sets (**Fig. 2D** and **Supplemental Fig. 4**). Chondrocyte-like osteoprogenitor (COP) cells,^31^ a subtype of metaphyseal progenitors, are also located within the chondrogenic clusters. Finally, we found that ACAR1-3 cell transcriptomes matched those of marrow adipogenic lineage precursors (MALP),^24^ skeletal stem and progenitor cells,^19^ flow sorted CAR cells,^32^ and stroma populations as defined by mass cytometry surface markers.^33^

We further confirmed stromal cell cluster hierarchies based on cell potency^34^ and lineage trajectories^35^. Consistent with our inferences from marker gene expression (**Fig. 2A-D**), MesenP1-2 cells showed the highest stemness potential scores (**Fig. 2E**). This was followed by the BiPot cluster, which possessed a wide potency distribution reflecting a progressive lineage restriction (**Fig. 2E**). Chondrolineage populations also showed varying degrees of pluripotency in these analyses, suggesting that these cells reflect partially committed progenitor cells rather than mature chondrocytes.

ACAR, OCAR, osteoblast (Ob) and osteocyte (Oc) cells were the most differentiated cell types by potency analysis (**Fig. 2E**). We next examined single lineage trajectories that gave rise to ACAR or osteolineage cells. We set *Hic^+^ Cd34^+^* MesenP1 cells as the earliest progenitors and ascertained an ACAR trajectory that traverses MesenP2 cells and BiPot cells (**Supplemental Fig. 5B**). While the relationship between MesenP1-2 requires further clarification, it is clear that MesenP cells become BiPot cells prior to becoming ACAR cells (**Fig. 2F** and **Supplemental Fig. 5B**). Similar pseudotime analysis experiments revealed a discrete trajectory from MesenP to BiPot to OCAR and Ob cells (**Fig. 2G** and **Supplemental Fig. 5C-D**). We were also able to recapitulate these trajectories upon aggregated analysis of all osteo- and ACAR lineages (**Supplemental Fig. 5E-G)**.

Taken together, these findings identify *bona fide* CAR cells based on transcriptional profiles and characterize the differentiation pathway of hematopoietic niche stroma from mesenchymal progenitors.

### The mouse microenvironment that supports hematopoiesis emerges in postnatal life

We next sought to understand the emergence of stromal and mesenchymal populations across the mouse life span. We stratified our atlas into five stages (fetal, neonatal, juvenile, adult, and aged) and observed dramatic shifts in cell types over development (**Fig. 3A**). We noted an expected predominance of mesenchymal and chondrolineage progenitor cells during fetal and neonatal periods, after which there was an increase in osteolineage cells (**Fig. 3B**).

**Figure 3.**
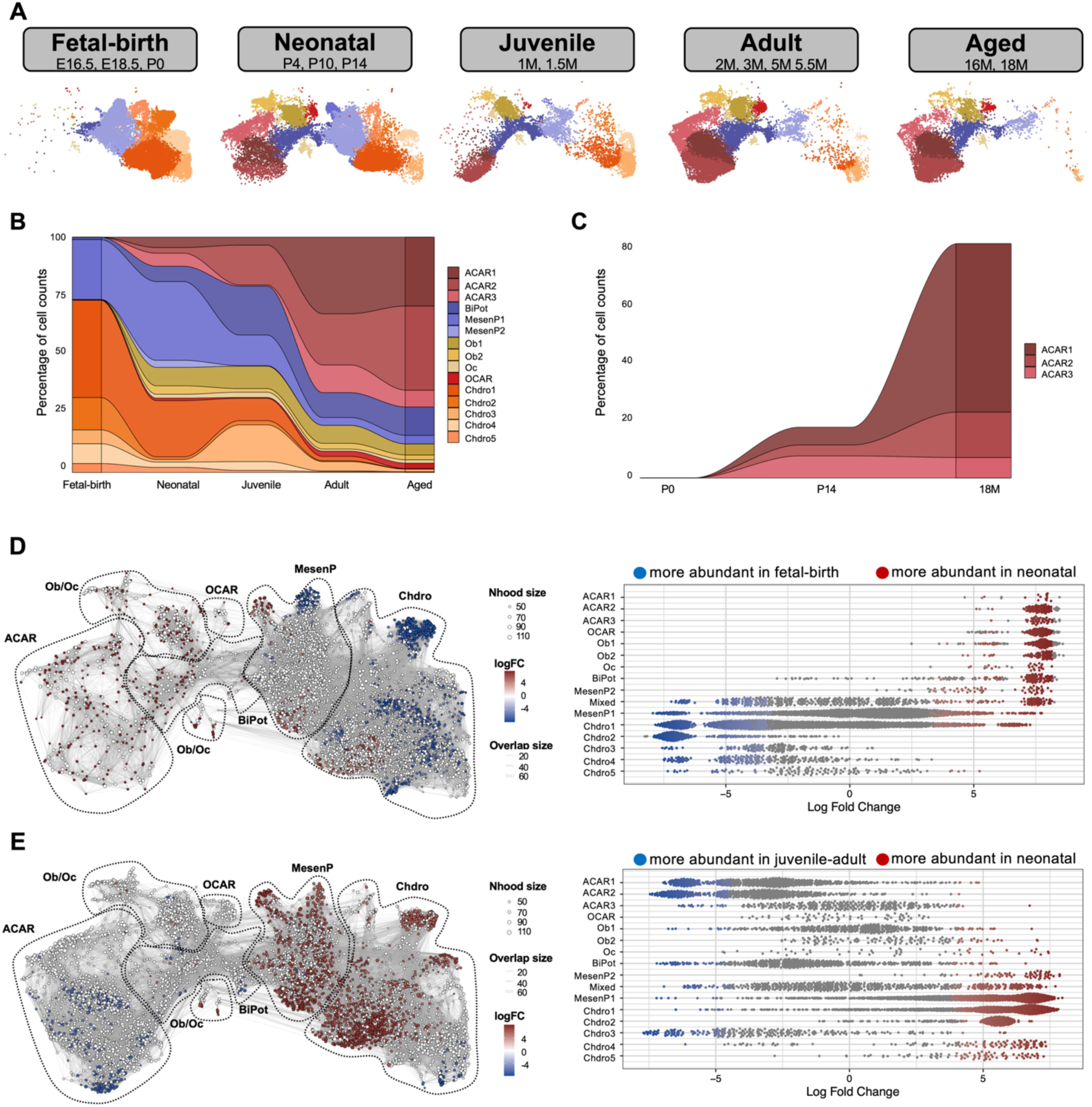
ACAR differentiation is blocked in fetal life. (A) UMAP projections of mesenchymal lineage cells divided by developmental stage. (B) Alluvial plot displaying the percentage of mesenchymal lineage clusters across development. (C) Alluvial plot displaying the percentage of ACAR cluster across development. (D-E) Differential abundance testing of cell states using MiloR comparing (D) fetal-birth and neonatal stages and (D) juvenile-adult and neonatal stages. Differential abundance testing is shown as a UMAP projection and beeswarm plot with significantly different groups of transcriptionally similar cells (e.g. neighborhoods) colored by magnitude and directionality of fold difference.

Hematopoietic niche-supporting ACAR cells were rare from fetal life through birth (E16.5-P0), but undergo significant rapid expansion in the postnatal bone marrow (**Fig. 3B-C**). Differentiated BiPot progenitors and ACAR were detected as early as P4, indicating that initial ACAR formation likely occurs soon after birth (**Supplemental Fig. 6A**).

We identified an apparent developmental ‘block’ in mesenchymal progenitor differentiation at the late MesenP1 and early BiPot progenitor stage during fetal life (**Figure 3A-C** and **Supplemental Fig. 6B**). To interrogate this maturation block, we performed differential abundance testing between different stages of mouse life with MiloR.^36^ When comparing fetal to neonatal time points, MesenP1 cells with highest pluripotency (**Fig. 2E**) were enriched during fetal life, whereas more lineage-committed BiPot cells were enriched in neonatal bone marrow (**Fig. 3D**). Comparison of neonatal to juvenile-adult marrow revealed a similar shift from uncommitted mesenchymal and chondrolineage cells in neonates to more differentiated BiPot and ACAR cells in the juvenile and adult stages (**Fig. 3E**). These findings place the emergence of mouse hematopoietic niche-supporting cells in the time just after birth.

We next sought to identify developmental stage-specific expression patterns in hematopoietic niche factors, adipokines, and osteokines over time. Factors associated with embryonic skeletal development (*Dlk1* and *Igf2*), and bone marrow regeneration^8^ and fetal bone marrow signaling (*Ptn*), were upregulated in the early postnatal period (**Fig. 4A**, **Supplemental Fig. 7**, and **Supplemental Table 3**). *Dlk1* expression predominates in fetal life and inhibits adipocyte differentiation.^37,38^ Postnatal decline in *Dlk1* levels could also thus facilitate CAR cell development, as this is an adipogenic lineage. *Col3a1*, which marks mouse fetal bone marrow stroma,^16^ is also highly expressed during the fetal and juvenile periods, supporting the accurate characterization of mesenchymal progenitors and marrow stroma in our data set.

**Figure 4.**
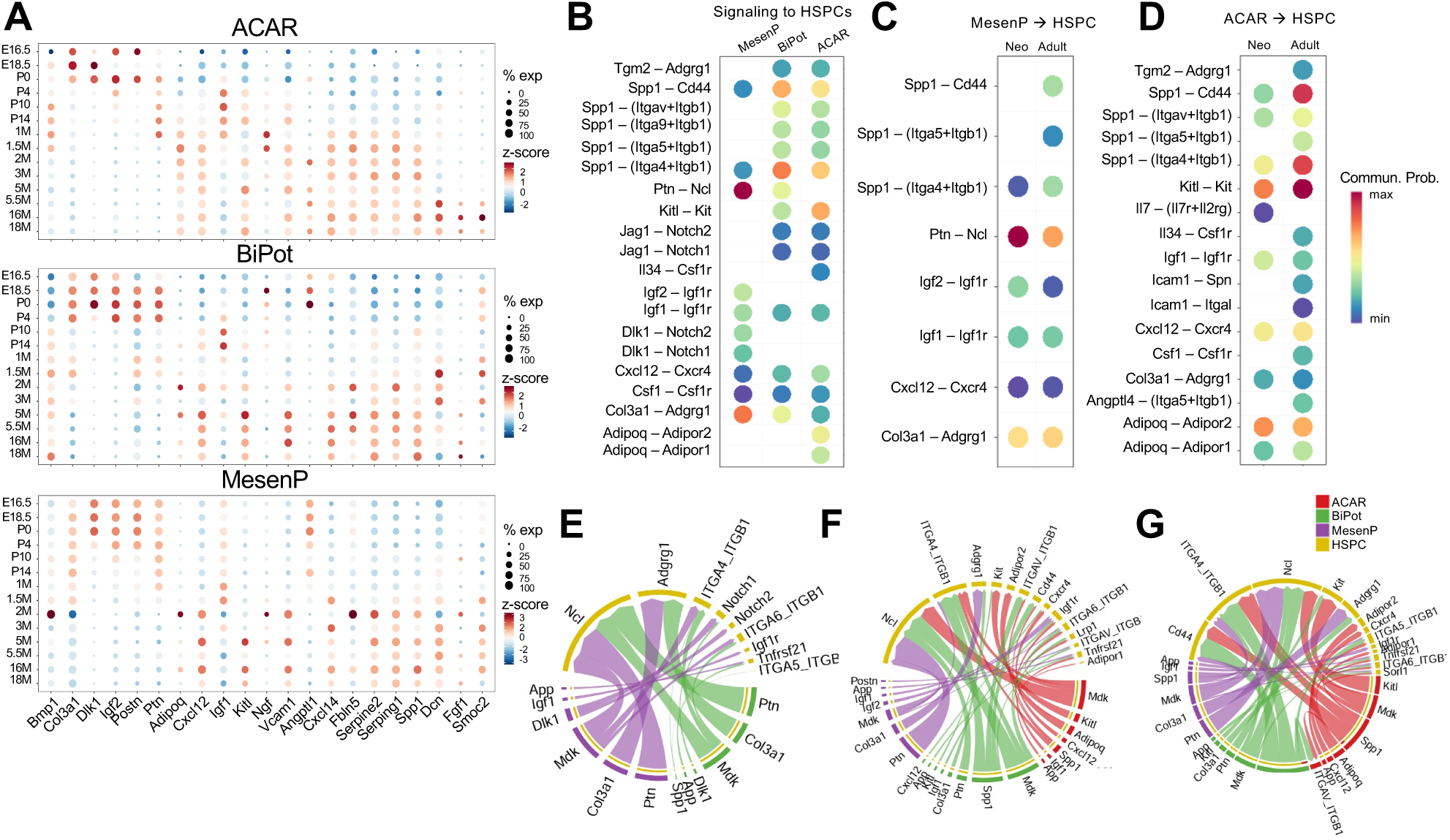
Hematopoiesis supporting secretome emerges in the postnatal bone marrow. (A) Dot plot of the expression of select niche factors, adipokines, or osteokines across development for indicated mesenchymal populations. ACAR and MesenP expression analysis represent pooled expression of all clusters. (B-G) Signaling between HSPCs and ACAR, BiPot, and MesenP cells was assessed using CellChat. (B-D) Dot plots of communication probability of outgoing signals from select pathways independent of developmental stage (E-F) Chord plots of cell-cell communication between mesenchymal populations and HSPCs by developmental time point: (E) Fetal-birth, (F) Neonatal, (G) Juvenile-adult stages.

Substantive expression of the HSC-supporting signals *Cxcl12*, *Kitl*, and *Igf1* first occurs in the neonatal period. These signals then persist through the marrow lifespan, although *Igf1* is somewhat downregulated in aged samples (**Fig. 4A**). This *Igf1* finding is consistent with prior work.^10^ Neonatal ACAR cells also upregulate *Ngf*, which supports marrow innervation.^39^ While most cytokines and matrix components persisted throughout adult time points, some genes were specifically associated with aged marrow (*Dcn, Smoc2, Fgf1*).

Analyses of active cell-cell communication in the bone marrow niche also supported postnatal emergence of functional ACAR-replete hematopoietic niche. Cxcl12-Cxcr4, Kitl-Kit, and Spp1-based signaling modalities were all enriched in ACAR cells compared to MesenP and BiPot cells (**Fig. 4B**), with MesenP cells providing some Osteopontin and Igf1/2 signaling to HSPCs in the neonatal and adult time periods (**Fig. 4C**). As expected, ACAR cells provide the most robust niche-supporting signals (e.g., Kitl, Cxcl12, Igf) and the strength of these signals increases from neonatal to adult life (**Fig. 4D**). Composite analyses of stromal-HSPC signaling at fetal, neonatal, and adult time points also supported the postnatal maturation of key signaling modalities including Cxcl12 and Kitl from ACAR cells (**Fig. 4E-G**).

These findings support a temporal model wherein hematopoietic niche cells create a hematopoietic-supportive bone marrow microenvironment in the neonatal period. Interestingly, it also appears that mesenchymal progenitor differentiation into ACAR cells is blocked in fetal life. This observation raised 2 key questions. 1) Do progressive transcriptional changes occur to support the development and maturation of niche-supporting stroma (e.g., ACAR cells)? 2) What mechanisms are responsible for alleviating the developmental block on mesenchymal progenitor cell differentiation in fetal life?

### Transcription factor activity is temporally regulated in the bone marrow niche

Our data suggest ACAR cells originate from MesenP cells (**Fig. 2F**). When we compared gene regulatory networks to infer transcription factor activities in MesenP and ACAR cells using SCENIC^40^, we found enrichment of relevant transcriptional regulators of limb mesenchyme and embryogenesis in MesenP (*Meox2, Sox11*, *Osr2*; **Fig. 5A**). In contrast, key transcription factors for CAR cells and adipolineage cells were increased in ACAR clusters (*Ebf3*, *Foxc1*, and *Pparg*; **Fig. 5A**). These transcriptional activities matched well with the RNA transcript abundance of these key factors, including *Ebf3* (**Fig. 5B** and **Supplemental Fig. 8**).

**Figure 5.**
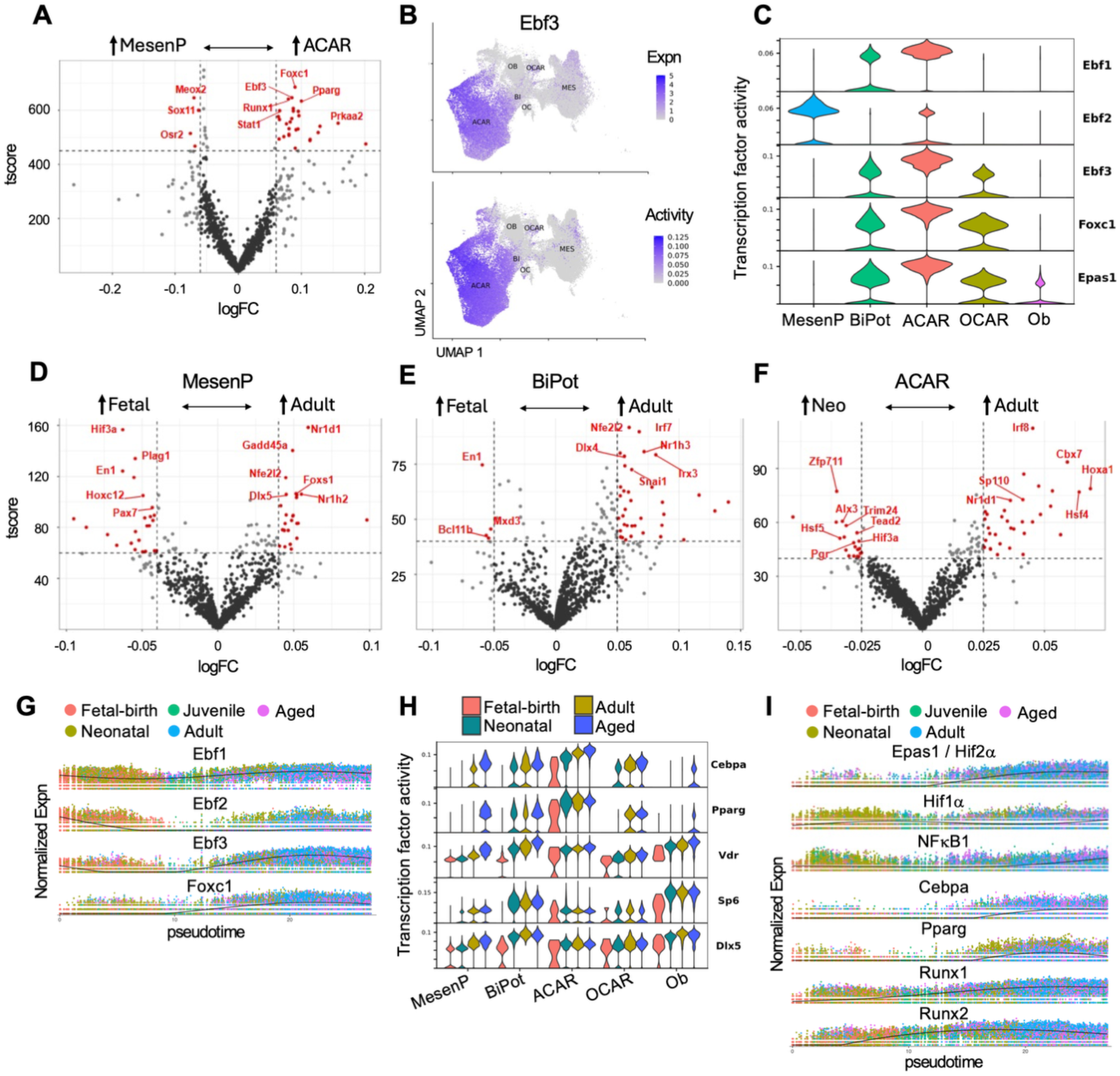
Transcription factor activity is developmentally controlled in the bone marrow niche. (A) Volcano plot of differential transcriptional activities comparing MesenP and ACAR cells independent of developmental time. (B) Feature plots of *Ebf3* expression (top) and activity (bottom). Transcription factor expression and inferred activity was assessed by differential gene expression and gene regulatory networks using Seurat and SCENIC, respectively. (C) Violin plot of key CAR cell transcription factor by adipo-osteolineage cluster. (D-E) Volcano plots of differential transcriptional activities by developmental stage in indicated mesenchymal progenitor or ACAR cells. (G) Key CAR cell transcription factor expression plotted across pseudotime using Monocle 3. Individual cells are colored by developmental stage. (H) Violin plots of select transcription factor activities in mesenchymal lineage populations grouped by developmental stage. (I) Adipo-osteolineage, HIF, and NfκB1 transcription factor expression plotted across pseudotime using Monocle 3. Individual cells are colored by developmental stage. For all volcano plots, select statistically different transcription factors are labeled red.

We then extended this analysis to other adipo-osteolineage factors and observed dramatic differences among Ebf activities. Ebf2 activity is high in MesenP cells but absent in BiPot cells, suggesting that *Ebf2* activities may restrict lineage commitment (**Fig. 5C**). We also identified a progressive increase in Ebf1, Ebf3, and Foxc1 activities along the MesenP-BiPot-ACAR developmental trajectory, which is starkly reduced upon osteolineage commitment (**Fig. 5C**).

To ascertain temporal transcription factor regulation, we compared transcriptional activities across cell types and over development. Transcription factors involved in the epithelial-mesenchymal transition (Snai1), inflammatory responses (Irf7, Irf8), and mesenchymal lineage commitment were dynamically regulated in the MesenP, BiPot, and/or ACAR populations (**Fig. 5D-F**). This prompted us to ask what pioneer factors and/or cell-extrinsic forces drive these changes in transcriptional activities.

We plotted CAR and adipo-osteolineage transcription factor expression over pseudotime to define molecules subject to temporal regulation. As expected, CAR cell factors emerge after birth (*Ebf3* and *Foxc1,* **Fig. 5G**). By comparison, *Ebf2* expression is extinguished in the early neonatal period, consistent with its putative role in preventing MesenP differentiation (**Fig. 5G**). Adipo-lineage regulators were minimally expressed from E16.5-P0 but exhibited rapid induction during the neonatal period (*Pparg* and *Cebpa*, **Fig. 5H**). Maximal activities of adipogenic transcription factors occurred in BiPot and ACAR cells at the juvenile-adult stages, and MesenP cells also showed adipogenic transcription factor activities in in aged marrow. Osteolineage transcription factors changed to a smaller degree across development (*Runx2*, *Dlx5*, *Vdr, Sp7,* **Fig. 5H-I**).

Taken together, these findings suggest that transcription factor activities, including key CAR cell transcription factors, are dynamically regulated in stromal and mesenchymal cells over bone marrow ontogeny. ‘Pioneer’ factors like *Ebf1*, *Runx2*, and *Dlx5* may prompt undifferentiated MesenP cells to adopt adipo-osteogenic fates.

### Systemic changes promote hematopoietic niche emergence in neonatal bone marrow

We also identified dynamic transcriptional regulatory signatures closely tied to cell-extrinsic stimuli, including hypoxia (*Hif1a*, *Hif2a*) and inflammation (*NFkB1,* **Fig. 5I)**. This led us to suspect that environmental factors might be linked to the developmental block in mesenchymal differentiation in perinatal time periods, and perhaps instigate or drive niche development. Pathway analyses comparing MesenP or BiPot cells in the fetal-to-neonatal transition revealed striking enrichment of inflammatory signaling processes (**Fig. 6A**). Similarly, ACAR cells showed an upregulation in TNFα and interferon responses in adults compared to neonatal bone marrow (**Fig. 6A**). This suggests that late gestation interferon pulses, which impact neonatal HSCs,^41^ might also underlie stromal cell emergence and/or development.

**Figure 6.**
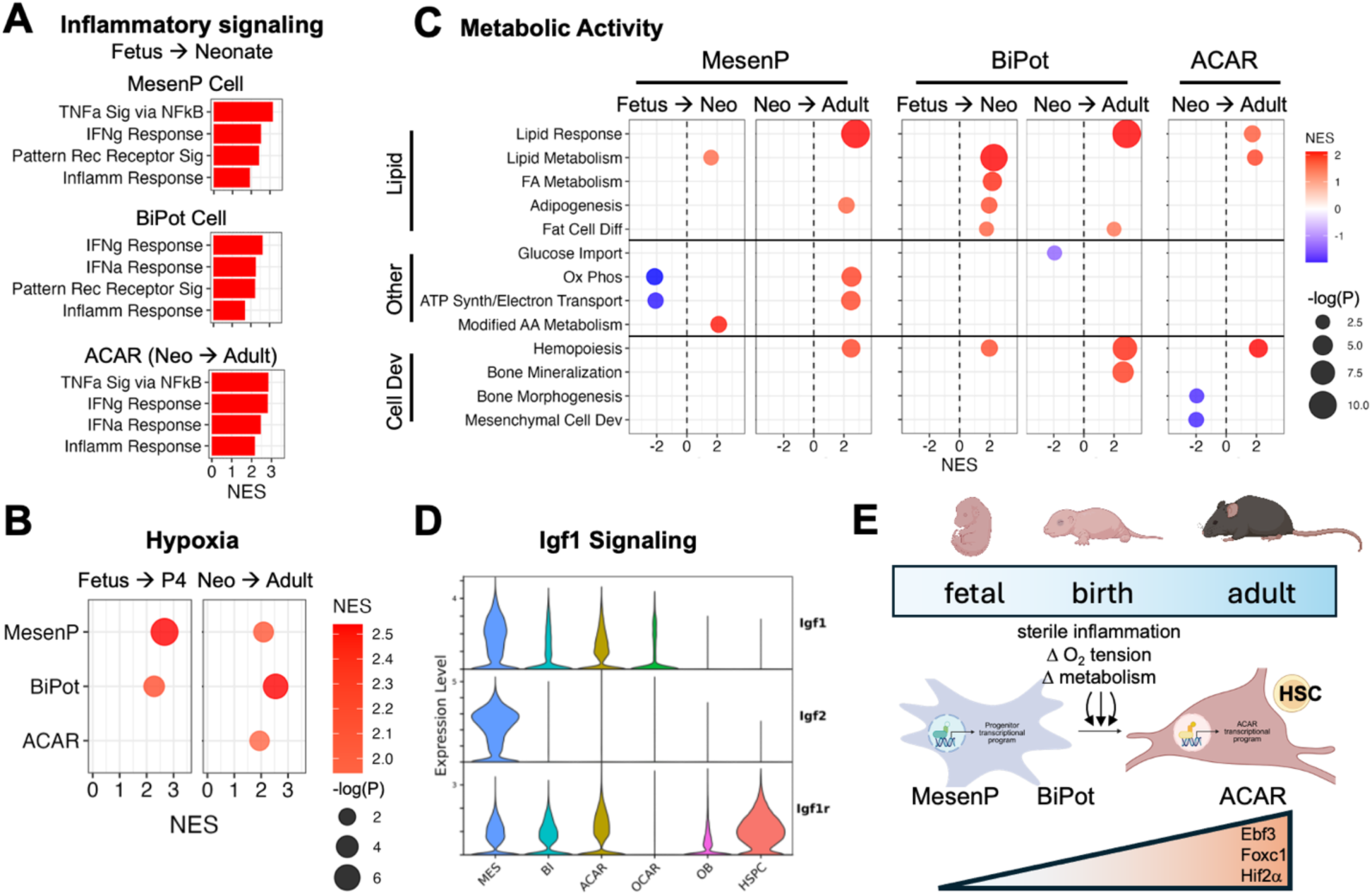
Transcriptional programs related to the fetal-to-postnatal transcription drive hematopoietic niche development postnatally. (A-C) Differential gene expression analysis followed by GSEA was performed on mesenchymal progenitor and ACAR populations across developmental stage. ACAR and MesenP analyses represent pooled expression of all clusters. Selected significantly enriched pathways are shown. (A) Bar plot of inflammatory pathway gene enrichment. (B) Dot plot of hypoxia gene enrichment. (C) Dot plot of metabolic pathway gene enrichment. Circle sizes represent conversions of p-values after multiple comparisons adjustment. (D) Violin plot of IGF pathway component expression by mesenchymal lineage cluster. (E) Proposed model of hematopoietic niche emergence in the perinatal period.

By gene set enrichment analysis (GSEA), we also identified hypoxic response signaling changes during the perinatal transition (**Fig. 6B**). Hypoxic responses occurred as early as P4 in MesenP and BiPot cells, whereas ACAR cells (which are essentially absent in fetal life) exhibit hypoxia responses at 2 months of age. The hypoxia response signal persisted in all stromal cells over time, although MesenP possess relatively few differentially expressed genes after the neonatal period (**Supplemental Fig. 8**). These findings suggest that changes in oxygen tension could promote changes in cell state and/or transcriptional response to direct niche formation in neonatal life. Importantly, these changes coincide with *Epas1* (HIF2α) induction (**Fig. 5C** and **Fig. 5I**).

In addition to hypoxic and inflammatory signals, we found changes in key metabolic pathways that appear to drive timely niche formation. By GSEA comparing MesenP, BiPot, and ACAR cell populations across time points, lipid and fatty acid oxidation pathways predominated after birth (**Fig. 6C**). This increased lipid processing power coincided with a shift away from oxidative phosphorylation in early MesenP cells during the neonatal period. It is likely that active Igf1 signaling helps to promote energy storage and growth in this context (**Fig. 6D**).

Changes in inflammation, hypoxia, and metabolic activities temporally coincide with changes in differentiation programs, particularly the dramatic shift at birth that derepresses the apparent lineage block in fetal mesenchymal progenitors. An absence of active differentiation programs may help explain why ACAR differentiation appears blocked prenatally, at a time when there are no external physiologic cues (**Fig. 3**). BiPot cells acquire pro-adipogenic and osteogenic differentiation gene expression in the neonatal period, consistent with potential lineage fates (**Fig. 6C**). One exception is the pro-adipogenic gene signatures in aged MesenP cells, which may reflect adipocyte accumulation characteristic of aging bone marrow and/or marrow exhaustion (**Fig. 6C**).^24^

In sum, these findings show that responses to external physiologic stimuli occur in temporal relation to dynamic transcription programs to drive bone marrow mesenchymal and stromal cells differentiation, including the initial development and maturation of ACAR cells critical for hematopoietic support (**Fig. 6E**). Inflammation, metabolic switches, and hypoxia may be key signals that foster timely hematopoietic niche emergence and permit HSC engraftment.^9^

## Discussion

Our longitudinal single cell atlas provides developmental characterization of BMSC emergence over mouse bone marrow ontogeny. Cell trajectories and developmental progressions shown herein establish a framework to understand drivers of perinatal development, creating a bone marrow niche that supports lifelong hematopoiesis. We propose that changes in inflammatory, metabolic, and oxygen-sensing pathways drive dynamic transcription factor activities, underlying a rapid developmental maturation in BMSC differentiation and niche formation after birth (**Fig. 6E**). In this way, dramatic physiologic changes that accompany the birth process are critical for establishing the marrow niche. We anticipate that our atlas will support further discoveries related to developmentally regulated changes in bone marrow maturation throughout the mouse lifespan.

Inflammatory, metabolic, and hypoxia-related pathway changes in mesenchymal progenitors appear to drive CAR cell emergence. These same signaling modalities impact HSC biology. For example, sterile inflammation is critical for HSC regulation, neutrophil emergence, and granulopoiesis in the neonatal period.^41–44^ Metabolic changes have also been shown to modulate HSC quiescence and developmental programs, and hypoxia is a hallmark of the hematopoietic niche and bone marrow.^45,46^ Thus, our findings provide parallel molecular targets and signaling processes that impact mesenchymal niche development, which are necessary for long-term HSC support.

Our results show that an absence of mature niches in fetal and early neonatal life is due the absence of true ACAR cells, rather than decreased function. An absence of ACAR cells corresponds to decreased secreted Cxcl12, Scf, and Igf1 signals, which are absent in fetal life and arise in the neonatal bone marrow. An absence of these secreted signals agrees with prior results.^9,15^ Importantly, we identify a lineage trajectory between fetal mesenchymal progenitor cells to ACAR populations to explain relevant developmental origins of these critical cell types. During late gestation, *Hic1*^+^*Cd34*^+^ progenitors^29^ represent the top of an adipo-osteogenic developmental hierarchy. Mesenchymal progenitors with transcriptional profiles reminiscent of EMPs^24^ and universal fibroblasts^47^ lie directly downstream of *Hic1^+^* progenitors. These spawn *Runx2^+^* progenitors that ultimately give rise to BiPot progenitors and terminal ACAR and osteolineage cells. Inhibition of differentiation occurs at the level of *Runx2^+^* progenitors, which are only released in postnatal life. Key cell types and/or cell states along the ACAR lineage trajectory warrant future investigation.

We observe heterogeneity in metabolic and hypoxic gene expression signatures across mesenchymal progenitor and ACAR cell populations. ACAR2 cells possess the greatest enrichment in hypoxia-related gene expression, whereas ACAR1 cells upregulate fatty acid metabolic pathways. It is conceivable that these differences relate to anatomic regionalization, as shown in adult bone.^48^ These findings compel future studies to ascertain the spatial orientation of ACAR cell populations with respect to hematopoietic niches over time.

Our findings add temporal resolution to transcriptional activities that correspond with niche emergence, beyond a need for *Ebf3* and *Foxc1* to drive CAR cell differentiation in the perinatal period. The balance between *Ebf2* and *Ebf3* activities appears critical. Upregulation of *Ebf2* in early mesenchymal progenitors is associated with a fetal progenitor-like program that represses CAR differentiation.^12,13^ Epas1 (HIF2α) activity also mirrors Foxc1 and Ebf3 activities, suggesting that hypoxia signaling acts in concert with key CAR transcription factors.

We also identify early transcriptional events in mesenchymal progenitors that may delineate CAR cell emergence. Pseudotime transcriptional changes in Runx1/2 activities suggest that these may act as pioneer factors to commit early mesenchymal progenitors to a CAR cell fate in fetal life. Consistent with this notion, conditional loss of either Runx1 and Runx2 in limb mesenchyme does not significantly affect hematopoietic niche development, but combined Runx1/Runx2 loss in *Ebf3*-expressing cells increased marrow fibrosis and reduced hematopoiesis.^49^ Overall, we interpret relevant transcriptional changes as molecular mediators linking perinatal physiology to niche development.

In sum, this longitudinal atlas lends sheds new light on bone marrow niche formation and function. Despite consistent stromal populations in human and mouse niches, HSCs engraft at different times during human and mouse development.^14,50,51^ The bone marrow of a full term mouse pup is more closely related to an extremely premature human infant born at ∼22-24 gestational age. This has developmental and translational implications related to perinatal pathology. Prematurity and perinatal illness affect >10% of all infants worldwide. Adverse perinatal events might impact long term HSC fitness and/or niche maintenance through disrupted inflammatory, metabolic, and/or hypoxic cues, predisposing to bone marrow disease and blood disorders. Much of our current understanding of bone marrow development has come from studies of adult samples at steady-state, stress, or during marrow reconstitution following myeloablation. While there appear to be conserved changes in pathways and signaling in perinatal bone marrow establishment and marrow reconstitution after myeloablation,^8,39^ our data emphasize unique advantages in studying the perinatal marrow that continues to mature after initial HSC colonization. Our results will propel future investigation into the cell types and processes that facilitate bone marrow niche construction to support lifelong hematopoiesis.

## Methods

### Bone preparation and processing

For BMSC isolation and scRNAseq at P10 and 18 mo, procedures are as previously described.^27^ Briefly, bone marrow cells were flushed out with cell culture media and collected from tibias and femurs after the epiphyseal ends were removed. The bone shaft was then cut into pieces and digested with STEMxyme1 (Worthington) and Dispase II (ThermoFisher Scientific). The digested cells and bone marrow cells were pooled together and passed through a 70 µm filter before being stained and FACS-sorted against hematopoietic and erythroid markers. BMSC were enriched as the 7-AAD^-^/Calcein AM^+^/CD71^-^/Ter119^-^/Lin (CD45/CD3/B220/CD19/Gr-1/CD11b)^-^ fraction.

### Single cell sequencing

scRNAseq libraries were constructed with the Single Cell 3’ Reagent Kits v3.1 (10x Genomics). Sequencing was performed with a NovaSeq sequencer using 100-cycle paired-end reads, generating 200 million reads per sample. Cell ranger (10x Genomics) was used for demultiplexing, extraction of cell barcode, and quantification of unique molecular identifiers (UMIs). Downstream analyses were performed as described below.

### Harmonization and integration

Single-cell RNA-seq datasets were obtained from the NCBI Sequence Read Archive (SRA) using the SRA Toolkit (v3.0.3). Sequencing reads were downloaded with the prefetch command and converted to FASTQ format using the fastq-dump with the --split-files option to generate read-pair files. FASTQ outputs were inspected to confirm proper formatting and expected read lengths. Associated metadata were compiled from the corresponding GEO entries or from information provided in the original publications. Raw sequencing reads were aligned to the *Mus musculus* (mm10) reference genome using Cell Ranger (v7.1.0; 10x Genomics) to generate gene–barcode UMI count matrices.

Count matrices were processed in Python (v3.12.11) using the scvi-tools package (v1.3.3). Minimal quality control filtering was applied to remove cells with fewer than 500 total UMIs and genes detected in fewer than 5 cells. Data integration and dimensionality reduction were performed using the SCVI probabilistic generative model, which learns per-cell library size and corrects batch effects across libraries in a shared latent space. Graph-based clustering was conducted using the Leiden algorithm, and low-dimensional visualization was performed using UMAP with a minimum distance parameter of 0.3. Latent embeddings were subsequently imported into R (v4.4.0) and Seurat (v5.3.0) for visualization, marker gene identification, and cell type annotation.

Harmony and scVI tools are two common integration strategies employed for atlas-level dataset integration. Since our single cell atlas is one of the first to attempt integration of multiple previously published datasets across development time, we sought to test both methods to determine which integration mitigated dataset-associated bias without affecting the underlying developmental biology. As a first step, we subjected each integration to k-nearest-neighbor batch effect test (kBET), integration local inverse Simpson’s Index (iLISI), and cell-type (cLISI) as measures of integration effectiveness.^52–54^ kBET testing showed poor mixing, which likely resulted from biologic differences related to development rather than batch effect related integration difficulties. Indeed, iLISI and cLISI analyses, alternative measures of integration adequacy, demonstrated that Harmony and scVI tools were both effective methods of integration for this type of atlas.

We next performed dimensionality reduction and cell clustering to understand which integration method produced the most biologically meaningful structure. We examined the distribution of key hematopoietic (*Gypa*, CD45, *Pf4*, *Lyz2*) and non-hematopoietic (*Col2a1*, *Pdgfra*, *Bglap*, *Cdh5*, *Lepr*) lineage markers genes, and demonstrated that scVI integration resulted in a greater separation of hematopoietic and non-hematopoietic clusters found in mouse bone marrow (**Supplemental Fig. 1**). As Harmony and scVI tools-based integrations possessed comparable computational measures of integration, but scVI tools appeared to maintain biological, particularly of the mesenchymal stroma clusters, we used scVI integrated object for downstream computational analyses.

### CellChat

To determine the intercellular signaling present within the mouse bone marrow dataset, as well as how the signaling differed between cell types and time points, we used CellChat V2.1.0. The input for CellChat is a Seurat object. Initially, we used the entire mouse bone marrow dataset as our input. We ran CellChat under default parameters according to the CellChat vignette “Full tutorial for CellChat analysis of a single dataset with detailed explanation of each function”. We then subdivided the Seurat object into 4 time points (Fetal Birth, Neonatal, Juvenile Adult, and Aged) and performed CellChat on each individual time point. To compare the cellchat objects for each timepoint, we also used the CellChat vignette “Full tutorial for comparison analysis of multiple datasets”. All Bubble Plots were generated using CellChat and all Violin Plots were generated using Seurat.

### SCENIC

We used pySCENIC to determine the inferred activity of various transcription factors in the mouse bone marrow dataset. In R V4.4.1, we removed all non-stromal cells from the full Mouse Bone Marrow Seurat object. The reduced Seurat object contained only Mesenchymal progenitors, Bipotential cells, Adipo-CARs, Osteo-CARs, Osteoblasts, and Osteocytes. Using the function Seu2Loom from SeuratExtend V1.1.0, the Seurat object was converted into a .loom file.

On a Linux HPC, we activated the python package pySCENIC V0.12.1 using Singularity V3.11.1. The raw counts matrix of the loom was used as the main input for pySCENIC. PySCENIC inferred the Gene Regulatory Networks of the transcription factors and their target genes. It then calculated the level of enrichment for each motif in each cell, before assigning an AUcell Score for each transcription factor and each cell. The final output of SCENIC was an AUC matrix. All scripts used derived from the official pySCENIC tutorial and were run under default conditions with the exception of the “no pruning” argument being set to true. The scripts used can be found in the supplemental materials.

For the purposes of visualization and comparison between cell types and time points, the AUC matrix was then added back into the Seurat object. Additionally, any AUC values below 0.05 were imputed to 0 to remove poorly enriched motifs. Violin Plots and Feature Plots were generated using Seurat V5.1.0, and Volcano Plots were generated using SeuratExtend.

### Trajectory Analysis

We computationally defined ACAR and Osteo-OCAR cell lineage trajectories using algorithms within the Monocle 3 package.^35^ To maintain the structure of mesenchymal lineage clusters, we ensured that the original reduction and clustering constructed using Leiden algorithm were used for clustering, cell ordering, and trajectory generation performed with Monocle. Trajectories for the ACAR cell lineage, osteo-OCAR cell lineage, and Adipo-osteolineage or trajectories by developmental time were constructed by subsetting clusters belonging to each lineage or developmental stage and subsequently performing Monocle pipeline. We also employed the *graph_test* and *plot_genes_in_pseudotime* functions to investigate transcription factor expression along mesenchymal lineage trajectories. We also calculated pluripotency and inferred trajectories with CellTrace2 using default settings.

### Differential Gene Expression and Gene Set Enrichment Analyses

We performed differential gene expression (DGE) analyses using the *FindMarkers* function in Seurat V5.1.0, which utilizes a non-parametric Wilcoxon rank sum test to identify differentially expressed genes. To assess gene expression differences as a function of developmental time, we pooled multiple time points per developmental stage (*e.g.* E16.5, E18.5, P0 vs P4, P10, P14 to compare fetal-birth and neonatal stages, respectively) for analyses. For populations where more than one cluster was identified including ACAR cells and mesenchymal progenitors, these were also pooled for DGE analyses. For all DGE analyses, only genes that were detected within at least 10% of cells were tested.

After DGE analyses we performed GSEA using the fgsea package.^55^ Genes with an adjusted p-value < 0.05 and a log_2_ fold change of > 0.5 were used to create gene lists for downstream enrichment analyses. The mouse hallmark and M5 ontology gene sets from the Molecular Signatures Database (MSigDB) were utilized.^56–58^ Bar and bubble plots were constructed using ggplot2.

### Differential abundance testing of cell states

Differential abundance testing of cell states grouped within transcriptionally similar neighborhoods using the MiloR package.^36^ We used the original reduction, principal components, and clustering derived from Leiden algorithm to maintain the structure of mesenchymal lineage clusters. Differential abundance testing was performed among fetal-birth, neonatal, and adult-juvenile developmental stages by grouping multiple time points per stage as described above. KNN graph was generated using k = 30 and d = 10. Differential abundance results were visualized using *plotNhoodGraphDA* and *plotDAbeeswarm* functions within miloR.

## Acknowledgements

This study was supported by the National Institutes of Health (K99 HL156052 to CST, R00 HL177827 to CST, T32 HL007150 to BMD, R01 AG077911 to FL), the American Academy of Pediatrics Marshall Klaus Perinatal Research Award (BMD), the Children’s Hospital of Philadelphia Skeletal Health Research Affinity Group Pilot Award (BMD/CST), and the Children’s Hospital of Philadelphia Foerderer Award (BMD).

## Data and Code Availability

New single cell RNA sequences data for P10 and 18 months are available on NCBI GEO under accession GSE318094 (reviewer token: **kxatiuikrhojdyh**). The code used to generate data and figures is available on Github (github.com/thomchr/2026.MouseStromaAtlas) and upon request.

