## Supplemental Information for "A single cell atlas defines perinatal factors that drive mouse bone marrow development"

**Supplementary Information**

### Supplemental Figures

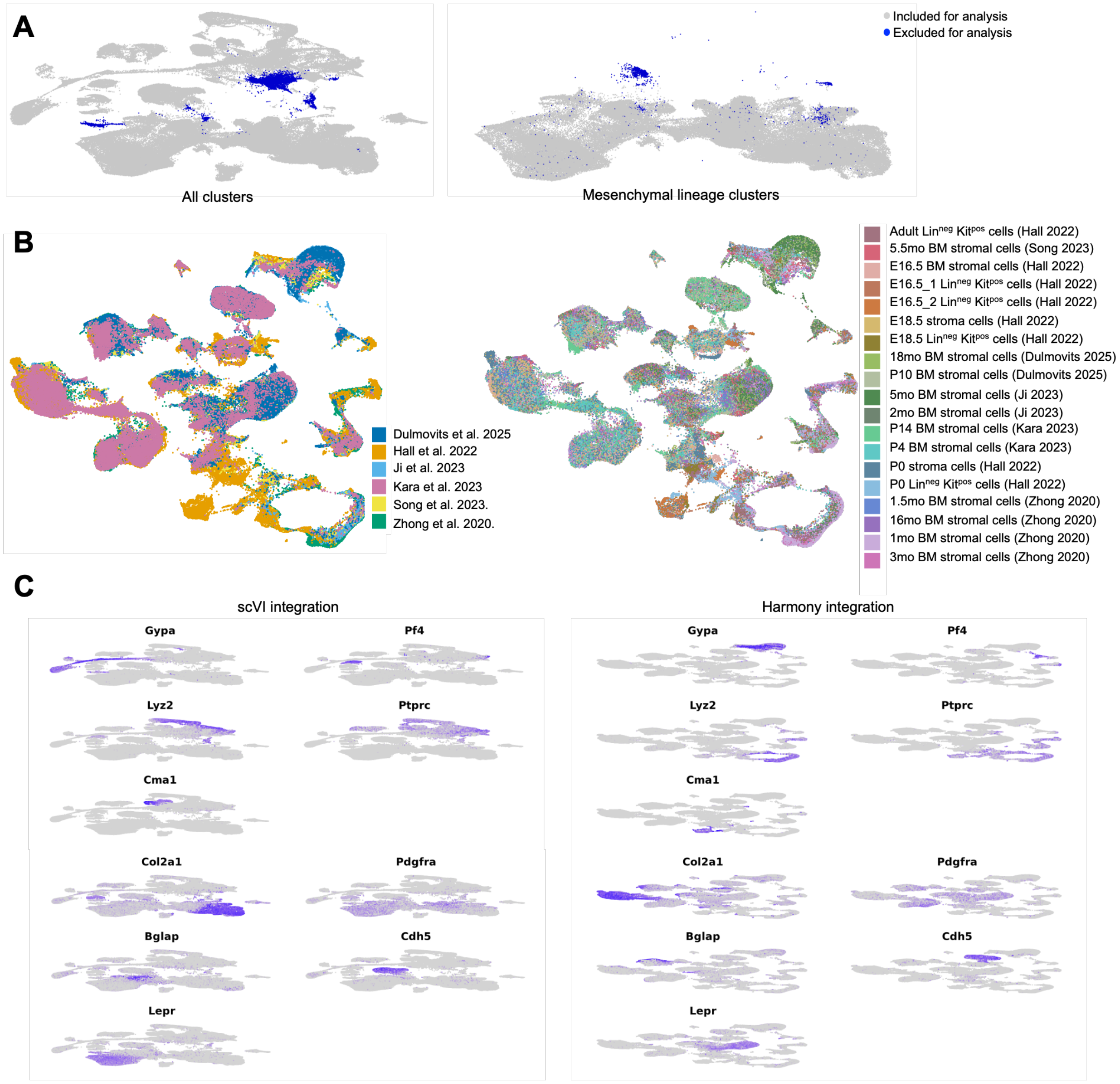


**Supplemental Figure 1. Quality control and comparison of integration methods.** (A) UMAP projections of complete dataset (left panel) and mesenchymal lineage clusters (right panel) with cells excluded for analysis highlighted. (B) UMAP projection of complete dataset using Harmony integration with individual cells colored by the dataset (left panel) and original scRNAseq sample (right panel). (C) Feature plots with select hematopoietic and stromal lineage markers using scVI or Harmony-based integration.

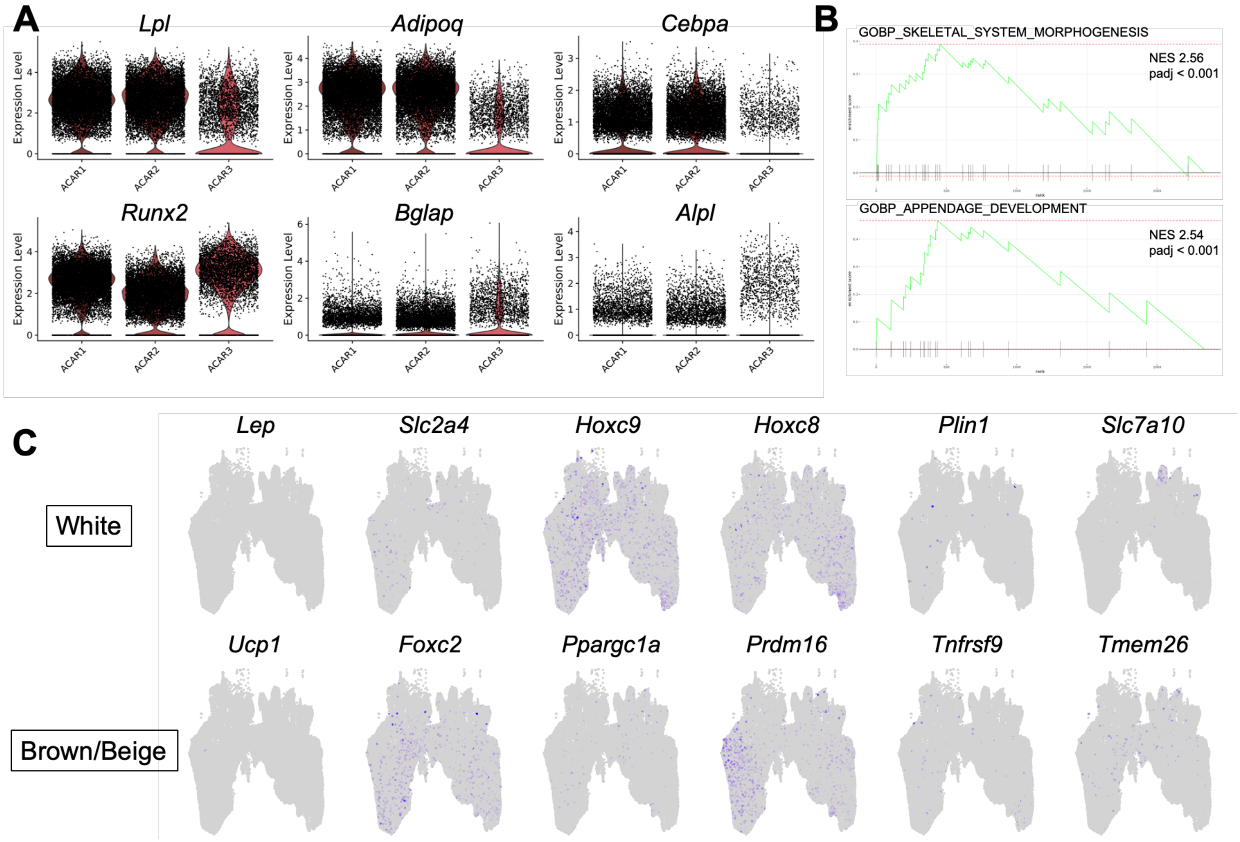


**Supplemental Figure 2. ACAR cells are transcriptionally heterogeneous and distinct from mature adipocytes.**
(A) Violin plot of the expression of select adipo- and osteolineage genes across ACAR clusters. (B) Differential gene expression and GSEA was performed by comparing ACAR3 to ACAR1 and ACAR2 clusters. GSEA plots of select statistically significant pathways demonstrating that ACAR3 cells possess an adipo-osteolineage transcriptional profile. (C) Feature plots with white, beige, and brown adipocyte marker genes demonstrating that most genes associated with different subtypes of mature adipocytes are lowly expressed in ACAR cells.

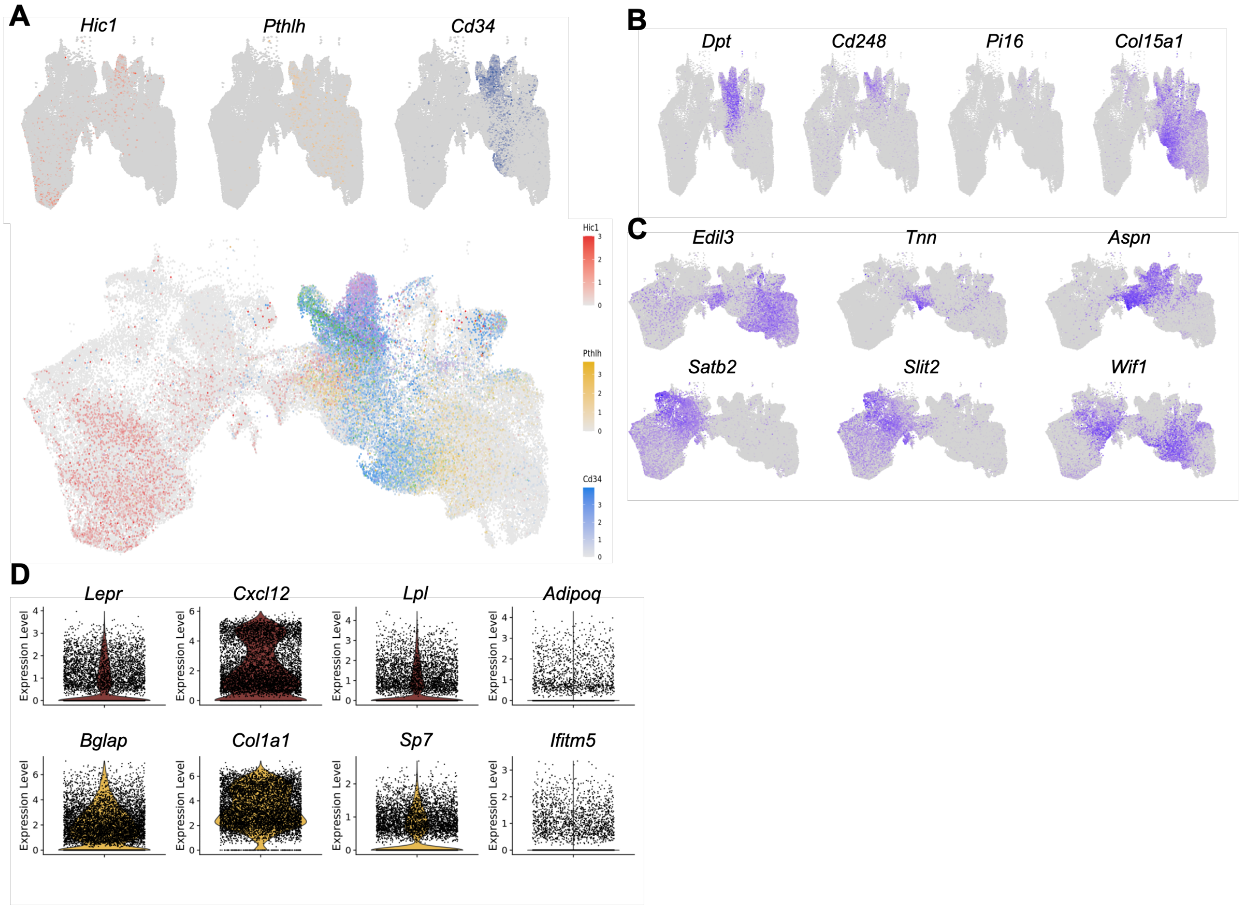


**Supplemental Figure 3. Single cell atlas identifies subpopulations of mesenchymal progenitors.**

(A) Individual feature plots of *Hic1*, *Pthlh*, *Cd34* (top panel) and combined feature plot with all three genes (bottom panel) marks *Hic1* and *Pthlh* expressing mesenchymal and skeletal stem cells. (B) Feature plots of select universal fibroblast marker genes suggesting that mesenchymal progenitors and fibroblasts possess overlapping gene expression profiles. (C) Feature plots of select marker genes demonstrate two subpopulations within the BiPot cluster. (D) Violin plot of the expression of select adipo- and osteolineage genes in BiPot cells.

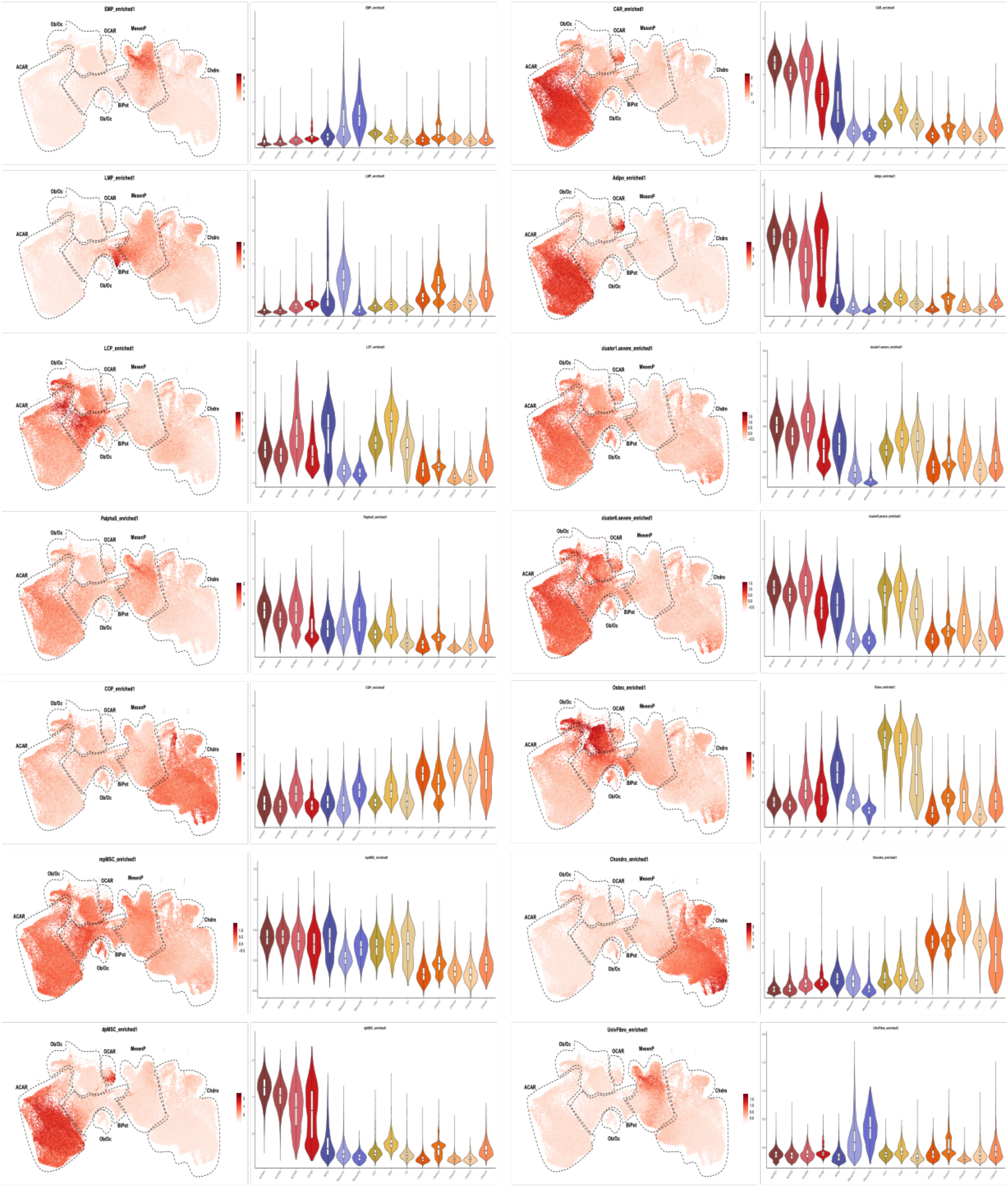


**Supplemental Figure 4. Single cell atlas contains previously characterized mesenchymal lineage populations.** Feature plot with corresponding violin plot of module scores related to previously published mesenchymal progenitor and terminal populations. EMP: early mesenchymal progenitor; LMP: late mesenchymal progenitor; LCP: lineage committed progenitor; P$\alpha$S: Pdgfr-alpha$^+$ Sca1$^+$ cell; COP: chondrocyte-like osteoprogenitor; mpMSC:

metaphyseal mesenchymal stromal cell; dpMSC: diaphysis mesenchymal stromal cell; CAR: Cxcl12 abundant reticular cell; Adipo: adipolineage; cluster1 and 6.severe: bone marrow stromal cells defined by cell surface marker mass cytometry. Osteo: Osteolineage; Chondro: chondrolineage; UnivFibro: universal fibroblast.

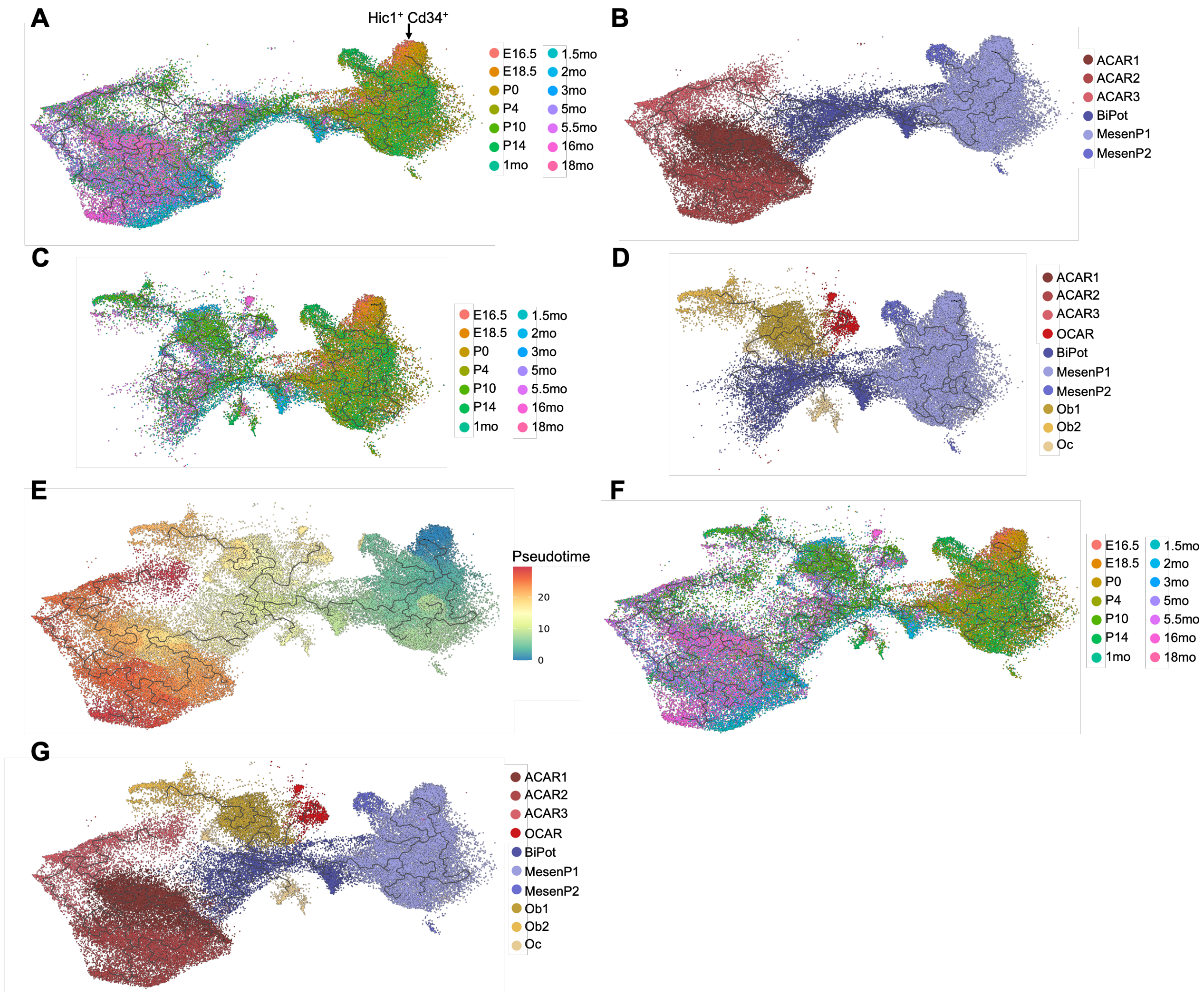


**Supplemental Figure 5. Defining the adipo-lineage trajectory. Lineage trajectories were constructed by arranging transcriptional cell states along pseudotime with Monocle3.** (A-B) ACAR cell lineage trajectory. (A) UMAP projection with superimposed lineage trajectory and individual cells colored by developmental time point. *Hic1*$^{+}$*Cd34*$^{+}$ cells predominantly derived from E16.5-P0 time points represent the earliest progenitors. (B) UMAP projection with superimposed lineage trajectory and individual cells colored by cluster. (C-D) Osteo-OCAR cell lineage trajectory. (C) UMAP projection with superimposed lineage trajectory and individual cells colored by developmental time point. (D) UMAP projection with superimposed lineage trajectory and individual cells colored by cluster. (E-G) Adipo-osteolineage trajectory. (E) UMAP projection with superimposed lineage trajectory and individual cells colored by pseudotime. ACAR and osteo-OCAR cell trajectories diverge at a shared bipotent progenitor. (F) UMAP projection with superimposed lineage trajectory and individual cells colored by developmental time point. (G) UMAP projection with superimposed lineage trajectory and individual cells colored by cluster.

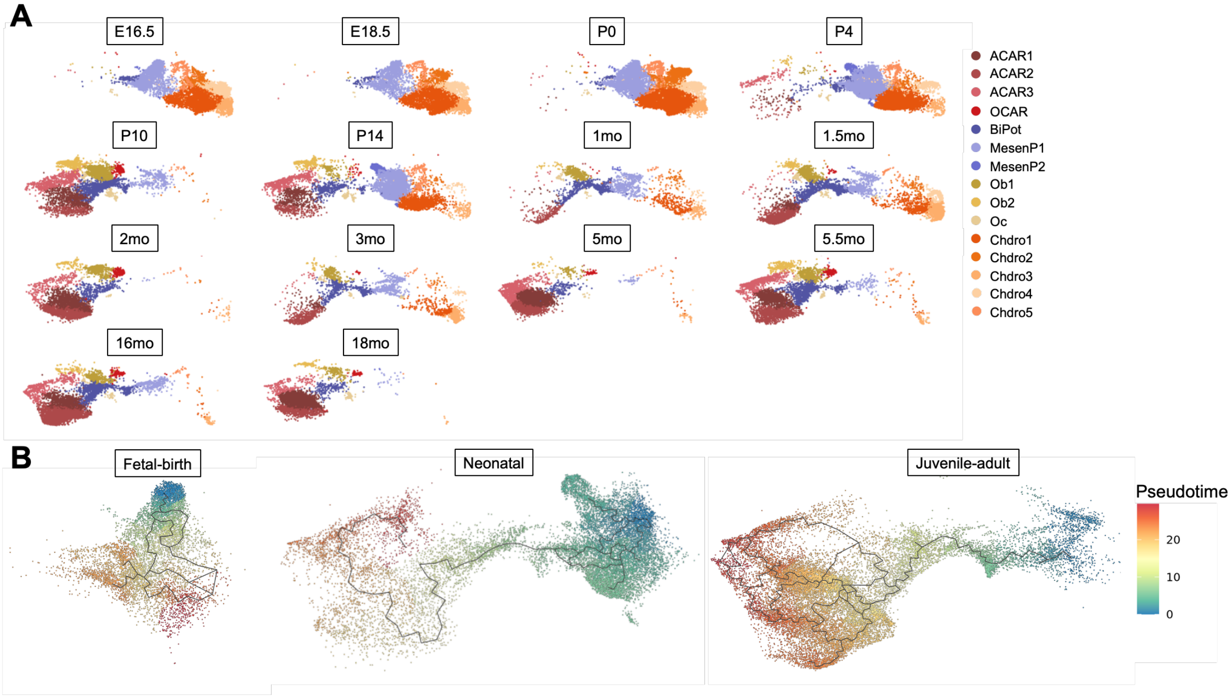


**Supplemental Figure 6. The absence of ACAR cells in fetal life is due to a block in mesenchymal progenitor differentiation.**
(A) UMAP projections of mesenchymal lineage clusters by developmental time point. ACAR cells are not observed until the postnatal period with the earliest detectable ACAR cells at P4.
(B) ACAR cell lineage trajectories were constructed by arranging transcriptional cell states along pseudotime with Monocle3 based on developmental time including fetal-birth, neonatal, and postnatal stages. The full ACAR lineage trajectory is not seen until after birth.

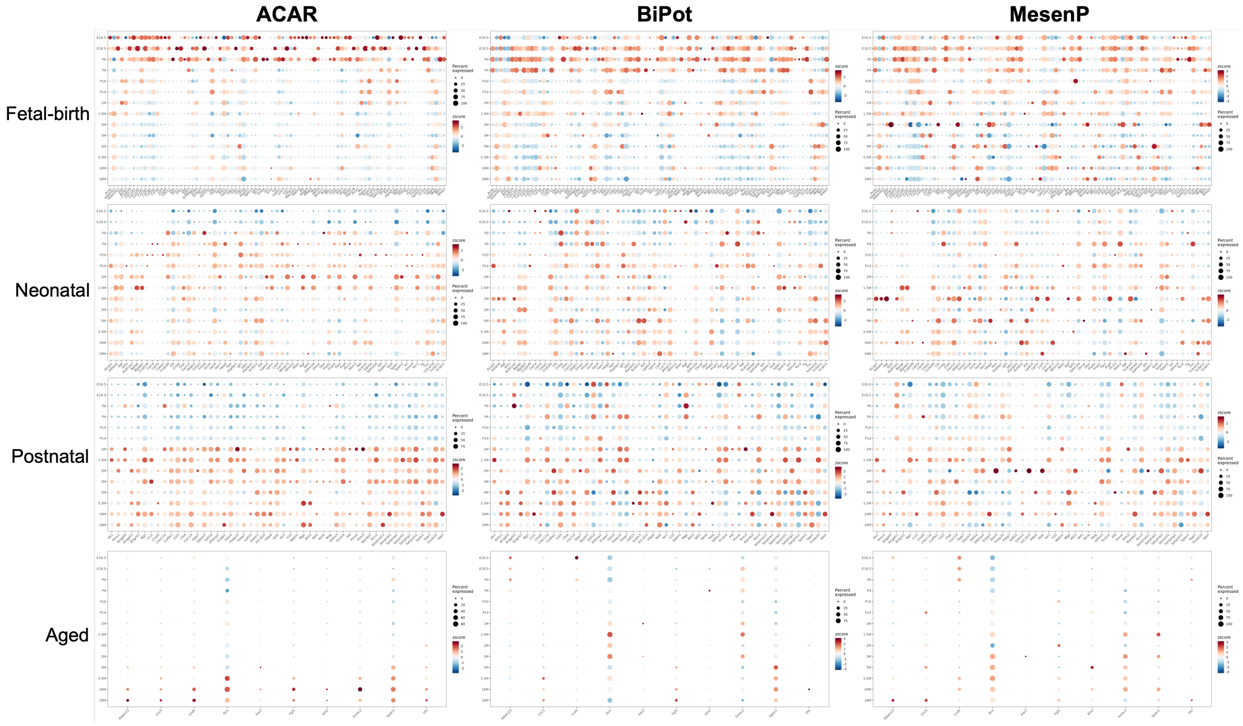


**Supplemental Figure 7. Profiling the ACAR and mesenchymal progenitor secretome across development.** Dotplots of the expression of niche-related factors, adipokines, and osteokines by developmental time point arranged into four groups (fetal-birth, neonatal, postnatal, and aged) based on when expression is observed. Plots are shown for pooled ACAR, BiPot, and pooled MesenP clusters.

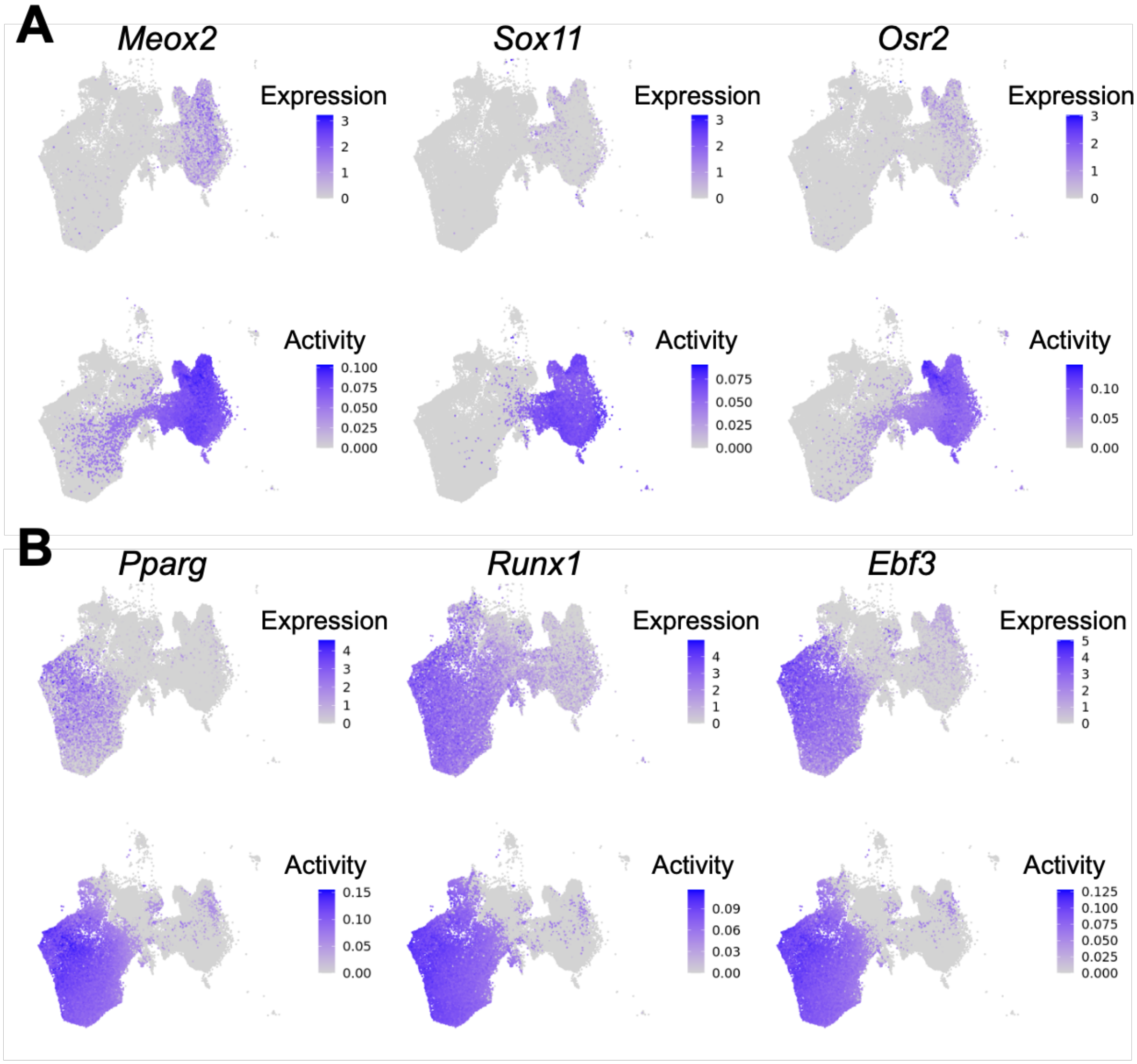


**Supplemental Figure 8. Mesenchymal lineage clusters demonstrate concordance with transcription factor expression and activity.**
(A-B) Feature plots of select transcription factor expression and activity for (A) mesenchymal progenitors and (B) ACAR cells.

**Supplemental Tables**

**Supplemental Table 1. Summary of datasets used to generate the single cell atlas of mouse bone marrow development.**

| **Dataset** Sample (# of mice) | **GEO/SRA accession** | **Bone source** | **Fraction isolated** | **Digestion** |
|---|---|---|---|---|
| Dulmovits | | | | |
| P10 BM stromal cells (3)** | GSE318094 | Femur Tibia | Lineage$^{neg}$, CD45$^{neg}$, CD71$^{neg}$ | Dispase II, STEMxyme1 |
| 18mo BM stromal cells (3)* | | | | |
| Hall *et al*. 2022 | | | | |
| *Hematopoietic cells* | | | | |
| Adult Lin$^{neg}$ Kit$^{pos}$ cells (4) | GSE178951 | Femur Tibia Pelvis Spine | Lineage$^{neg}$, CD45$^{pos}$, Kit$^{pos}$ | Collagenase II DNaseI |
| E16.5_1 Lin$^{neg}$ Kit$^{pos}$ cells (38)** | | | | |
| E16.5_2 Lin$^{neg}$ Kit$^{pos}$ cells (38)** | | | | |
| E18.5 Lin$^{neg}$ Kit$^{pos}$ cells (9)** | | | | |
| P0 Lin$^{neg}$ Kit$^{pos}$ cells (6)** | | | | |
| *Stromal cells* | | | | |
| E16.5 BM stromal cells (38)** | GSE178951 | Femur Tibia Pelvis Spine | CD45$^{neg}$, Ter119$^{neg}$ | Collagenase II DNaseI |
| E18.5 BM stromal cells (9)** | | | | |
| P0 BM stromal cells (6)** | | | | |
| Ji *et al*. 2023 | | | | |
| 2mo BM stromal cells (3) | GSE232738 | Femur Tibia | Lineage$^{neg}$, CD45$^{neg}$, CD71$^{neg}$ | Dispase II STEMxyme1 |
| 5mo BM stromal cells (3) | | | | |
| Kara *et al*. 2023 | | | | |
| P4 BM stromal cells (n/a)** | PRJNA835050 | Femur Tibia | Lineage$^{neg}$, CD45$^{neg}$, CD140$^{pos}$ | DNaseI Liberase$^{DL}$ |
| P14 BM stromal cells (n/a) | | | | |
| Song *et al*. 2023 | | | | |
| 5.5mo BM stromal cells (3) | GSE221936 | Femur Tibia | Lineage$^{neg}$, CD45$^{neg}$, CD71$^{neg}$ | Dispase II STEMxyme1 |
| Zhong *et al*. 2020 | | | | |
| 1mo BM stromal cells (2) | GSE145477 | Femur Tibia | *Col2a1-Cre* TdTomato$^{pos}$ | Collagenase A Trypsin |
| 1.5mo BM stromal cells (3) | | | | |
| 3mo BM stromal cells (3) | | | | |
| 16mo BM stromal cells (3) | | | | |

Unless specified, bone marrow cells for single cell sequencing were derived from female and male mice. * All cells derived from male mice. ** For embryonic and neonatal samples, sex was not determined.

**Supplemental Table 2. Summary of mouse strains used to generate the single cell atlas of mouse bone marrow development.**

| **Dataset**<br>Sample | **Mouse strain (Identifier)** |
|---|---|
| Dulmovits | |
| P10 BM stromal cells | C57BL/6J (RRID:IMSR_JAX:000664) |
| 18mo BM stromal cells | |
| Hall *et al*. 2022 | |
| *Hematopoietic cells* | |
| Adult $Lin^{neg}$ $Kit^{pos}$ cells | C57BL/6J (RRID:IMSR_JAX:000664) |
| E16.5_1 $Lin^{neg}$ $Kit^{pos}$ cells | |
| E16.5_2 $Lin^{neg}$ $Kit^{pos}$ cells | |
| E18.5 $Lin^{neg}$ $Kit^{pos}$ cells | |
| P0 $Lin^{neg}$ $Kit^{pos}$ cells | |
| *Stromal cells* | |
| E16.5 BM stromal cells | C57BL/6J (RRID:IMSR_JAX:000664) |
| E18.5 BM stromal cells | |
| P0 BM stromal cells | |
| Ji *et al*. 2023 | |
| 2mo BM stromal cells | C57BL/6J (RRID:IMSR_JAX:000664) |
| 5mo BM stromal cells | |
| Kara *et al*. 2023 | |
| P4 BM stromal cells | C57BL/Ka (MGI: 2159822) |
| P14 BM stromal cells | |
| Song *et al*. 2023 | |
| 5.5mo BM stromal cells | C57BL/6J (RRID:IMSR_JAX:000664) |
| Zhong *et al*. 2020 | |
| 1mo BM stromal cells | *Col2a1-Cre* (RRID:IMSR_JAX: 003554)<br>*Rosa26* TdT (RRID:IMSR_JAX: 007914) |
| 1.5mo BM stromal cells | |
| 3mo BM stromal cells | |
| 16mo BM stromal cells | |

**Supplemental Table 3. Bone marrow secretome gene lists**

| | Genes |
|---|---|
| Niche factors | *Cxcl12*, *Kitl*, *Angptl1*, *Igf1*, *Ptn*, *Spp1*, *Pf4*, *Tgfb1*, *Jag1*, *Vcam1*, *Fgf1*, *Csf3*, *Ackr1*, *Thpo* |
| Adipokines | *Cxcl12*, *Adipoq*, *Lep*, *Retn*, *Rarres2*, *Grn*, *Rbp4*, *Ccn4*, *Fabp4*, *Serpine1*, *Fstl1*, *Ccl2*, *Sparc*, *Sparcl1*, *Itln1*, *Azgp1*, *Sfrp5*, *C1qtnf3*, *Lcn2*, *Serpina12*, *Il10*, *Il1a*, *Metrnl*, *Cxcl14*, *Il6*, *Igf1*, *Igf2*, *Fgf2*, *Mstn*, *Wnt10b*, *Gdf15*, *Ngf*, *S100a1*, *S100b*, *Nrg4*, *Vegfa*, *Vegfc*, *Bmp8b*, *Bmp7*, *Fgf21*, *Igfbp2*, *Edn1* |
| Osteokines (Liang *et al*. 2024) | *Acan*, *Acat2*, *Acp5*, *Acyp2*, *Adam23*, *Adamts2*, *Adamts3*, *Adamtsl1*, *Adgrb2*, *Adprhl1*, *Agt*, *Aif1l*, *Ak1*, *Aldh1a2*, *Aldh6a1*, *Alpl*, *Amy1*, *Angpt4*, *Angptl1*, *Angptl7*, *Ap2b1*, *Apoc1*, *Aqp4*, *Aspn*, *Atp1b2*, *B3gnt2*, *Bex1*, *Bglap*, *Bglap2*, *Bglap3*, *Bgn*, *Blvrb*, *Bmp1*, *Bpnt1*, *C1qa*, *C1qtnf9*, *Calca*, *Calcb*, *Cap2*, *Capg*, *Casq1*, *Casq2*, *Ccdc80*, *Cd109*, *Cd5l*, *Cela1*, *Cenpw*, *Ces2e*, *Cfd*, *Cfh*, *Cgref1*, *Chad*, *Chchd10*, *Chil3*, *Chst5*, *Cilp2*, *Cirbp*, *Ckm*, *Ckmt2*, *Clec11a*, *Clec3a*, *Clec3b*, *Clic5*, *Cntfr*, *Coch*, *Col10a1*, *Col11a1*, *Col11a2*, *Col12a1*, *Col1a1*, *Col1a2*, *Col2a1*, *Col3a1*, *Col5a1*, *Col5a2*, *Col6a1*, *Col8a1*, *Col9a1*, *Col9a3*, *Colgalt1*, *Colgalt2*, *Comp*, *Cp*, *Cpe*, *Cpz*, *Crabp2*, *Cryab*, *Cst3*, *Cthrc1*, *Ctrl*, *Ctsa*, *Ctsc*, *Ctsg*, *Ctsh*, *Ctsk*, *Cxcl12*, *Cxcl13*, *Dcn*, *Dctpp1*, *Ddah2*, *Def6*, *Des*, *Dkk1*, *Dlk1*, *Dmp1*, *Dnajb4*, *Dpp7*, *Dpt*, *Dpysl3*, *Ear2*, *Ecm2*, *Edil3*, *Eef1d*, *Efemp2*, *Egflam*, *Eif4ebp2*, *Eif4h*, *Elane*, *Enpp2*, *Entpd3*, *Epyc*, *F13a1*, *Fabp3*, *Fam20c*, *Fap*, *Fbln5*, *Fbln7*, *Fbp1*, *Fbp2*, *Fetub*, *Fgf1*, *Fkbp10*, *Fkbp7*, *Fmod*, *Frzb*, *Galnt1*, *Gas2*, *Gatm*, *Gclm*, *Gins1*, *Glb1*, *Glg1*, *Glrx3*, *Gm525*, *Gmppb*, *Gnas*, *Gpc1*, *Gpc6*, *Gpha2*, *Gpx7*, *Grem1*, *H2-Q10*, *H6pd*, *Hapln1*, *Hoxd4*, *Hpgd*, *Hpx*, *Hs3st3a1*, *Hsd17b11*, *Hspb6*, *Hspbp1*, *Ibsp*, *Ier2*, *Igdcc4*, *Igf1*, *Igfals*, *Igfbp4*, *Igkc*, *Iglc2*, *Igll1*, *Iglon5*, *Il17b*, *Il1r2*, *Inhba*, *Isl1*, *Itih2*, *Ivd*, *Jchain*, *Jdp2*, *Kazald1*, *Kctd12*, *Khk*, *Klk1*, *Klrb1a*, *Lect2*, *Lgals1*, *Lgals3*, *Lgi3*, *Lgmn*, *Loxl3*, *Lrp4*, *Ltbp3*, *Luc7l*, *Lum*, *Macrod1*, *Magix*, *Mapkapk3*, *Matn3*, *Mb*, *Mepe*, *Mettl21c*, *Mfap4*, *Mfap5*, *Mgp*, *Mia*, *Mill2*, *Mlf1*, *Mmp13*, *Mmp14*, *Mmp9*, *Mmrn2*, *Mpo*, *Mstn*, *Mthfs*, *Mustn1*, *Myl9*, *Myoc*, *Ncan*, *Ncmap*, *Nid2*, *Nmb*, *Nog*, *Npm1*, *Nr4a1*, *Nrg1*, *Ntf3*, *Ntn1*, *Ntrk1*, *Oit3*, *Olfml2b*, *Olfml3*, *Omd*, *Orm1*, *Oscar*, *Otor*, *P3h1*, *P3h3*, *P4ha1*, *Pawr*, *Pcdh17*, *Pcolce*, *Pdcd6*, *Pdlim3*, *Pf4*, *Pgam2*, *Pla2g5*, *Plod1*, *Plod2*, *Pmvk*, *Pnpt1*, *Postn*, *Pou6f1*, *Ppic*, *Ppp1r8*, *Ppp2r1b*, *Prg2*, *Prg4*, *Prok2*, *Prr15*, *Prrx1*, *Prtn3*, *Pthlh*, *Ran*, *Rbm3*, *Rbp4*, *Rcn1*, *Rcn3*, *Ret*, *Retnla*, *Robo1*, *Rpl35a*, *Rspo3*, *Rspo4*, *S100a1*, *S100a4*, *S100a5*, *S100b*, *Scara3*, *Scarf2*, *Sdcbp*, *Sec23a*, *Sema3d*, *Serpina1b*, *Serpina1d*, *Serpina3b*, *Serpina3f*, *Serpina3n*, *Serpind1*, *Serpine1*, *Serpine2*, *Serpinf1*, *Serping1*, *Serpinh1*, *Sfn*, *Siglec1*, *Slc38a10*, *Slco5a1*, *Slit3*, *Smoc2*, *Smpdl3a*, *Smtnl2*, *Snrpa*, *Sost*, *Sparc*, *Sparcl1*, *Spock2*, *Spon1*, *Spon2*, *Spp1*, *Srl*, *Srm*, *Stc2*, *Stmn4*, *Stub1*, *Syt8*, *Tcn2*, *Tg*, *Tgfb1*, *Tgfbi*, *Thbs2*, *Thbs3*, *Timm8a1*, *Tk1*, *Tmsb10*, *Tnc*, *Tnfrsf11b*, *Tnfrsf19*, *Tnfsf11*, *Tnn*, *Tnni1*, *Tnni2*, *Tnnt2*, *Tpd52l1*, *Tppp3*, *Trmt112*, *Ube2i*, *Ucma*, *Vasn*, *Vat1*, *Vcan*, *Vim*, *Vtn*, *Wif1*, *Wwp2* |
